# Overlapping binding sites underlie TF genomic occupancy

**DOI:** 10.1101/2024.03.05.583629

**Authors:** Shubham Khetan, Martha L. Bulyk

## Abstract

Sequence-specific DNA binding by transcription factors (TFs) is a crucial step in gene regulation. However, current high-throughput *in vitro* approaches cannot reliably detect lower affinity TF-DNA interactions, which play key roles in gene regulation. Here, we developed PADIT-seq (<u>p</u>rotein <u>a</u>ffinity to <u>D</u>NA by *in vitro* transcription and RNA <u>seq</u>uencing) to assay TF binding preferences to all 10-bp DNA sequences at far greater sensitivity than prior approaches. The expanded catalogs of low affinity DNA binding sites for the human TFs HOXD13 and EGR1 revealed that nucleotides flanking high affinity DNA binding sites create overlapping lower affinity sites that together modulate TF genomic occupancy *in vivo*. Formation of such extended recognition sequences stems from an inherent property of TF binding sites to interweave each other and expands the genomic sequence space for identifying noncoding variants that directly alter TF binding.

**One-Sentence Summary:** Overlapping DNA binding sites underlie TF genomic occupancy through their inherent propensity to interweave each other.

## Main Text

Regulation of gene expression in response to diverse intracellular and extracellular signals depends on sequence-specific DNA binding by transcription factors (TFs) to *cis*-regulatory elements. Hundreds of TFs have been profiled by high-throughput *in vitro* binding assays, such as protein binding microarrays (PBMs) (*1–6*), systematic evolution of ligands by exponential enrichment and sequencing (SELEX-seq) (*7*) HT-SELEX (*8*), Spec-seq (*9*), DAP-seq (*10*), SMILE-seq (*11*), and bacterial one-hybrid (B1H) assays (*12*). While *in vitro* data on TF DNA binding specificities have been powerful in helping to understand mechanisms underlying *in vivo* TF occupancy (*13–16*), nucleotides flanking motif matches in the genome have been found to alter TF occupancy (*14, 17–24*). While some flanking sequence effects have been shown to be due to particular DNA shape preferences of TFs (*14*), we hypothesized that gaps in the ability of *in vitro* TF DNA binding data to fully explain *in vivo* TF binding may be due to an inability to reliably detect lower affinity TF-DNA interactions.

Low affinity binding sites are important for precise spatial and temporal control of gene expression during development (*25–30*). Homotypic clustering of multiple, non-overlapping low affinity binding sites adjacent to each other can serve regulatory roles by increasing local TF concentrations, either cooperatively (*31*) or non-cooperatively (*32*), to achieve phenotypic robustness (*28, 31*). Multiple low affinity binding sites in short tandem repeats (STRs) have also been recently shown to increase local TF concentrations (*33*). Given their importance, high-throughput *in vitro* binding assays, such as BET-seq (*34*), HiP-FA (*35*), MITOMI (*36*), Spec-seq (*9*) and STAMMP (*37*), have been developed to quantify TF-DNA interaction strength and identify lower affinity binding sequences. However, these techniques require prior knowledge of a TF’s binding preferences to design the pool of sequences that are assayed. To address this gap and characterize additional mechanisms by which low affinity binding sites might play important roles in gene regulation, we have developed <u>p</u>rotein <u>a</u>ffinity to <u>D</u>NA by *in vitro* transcription and RNA <u>seq</u>uencing (PADIT-seq), a high-throughput technology to measure TF-DNA binding preferences at far greater sensitivity than prior high-throughput methods.

PADIT-seq is based on a synthetic genetic circuit whose output is directly proportional to TF-DNA binding affinity and specificity (Figure 1A). A constitutive T7 promoter drives *in vitro* transcription of a TF DNA binding domain (DBD) fused to an ALFA tag (*38*). Following *in vitro* translation, the DBD binds to candidate DNA binding sites and recruits T7 RNA Polymerase via an anti-ALFA nanobody, which increases the rate of downstream reporter gene expression in proportion to the strength of the TF-DNA interaction. In the absence of TF-DNA interaction, a minimal “D1” T7 promoter drives low levels of reporter gene expression (*39*). In each well of a 96-well plate, a highly diverse library of TF binding sites (TFBS) is assayed for binding by a particular DBD. Barcodes (BCs) in the reporter gene identify the TFBS that were bound. The same reporter plasmid library is used to interrogate the binding preferences of different DBDs of interest in separate wells.

**Figure 1.**
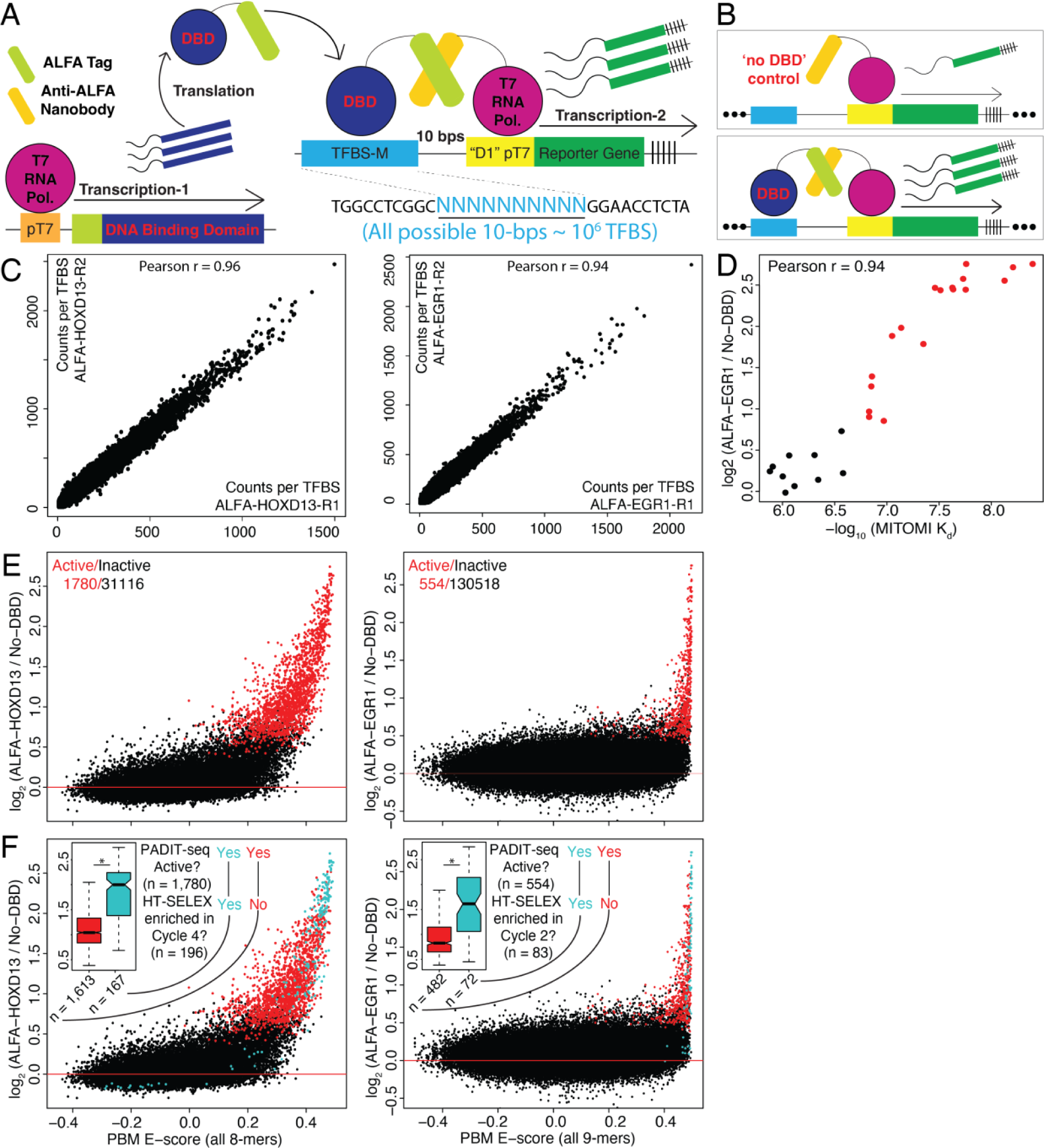
PADIT-seq detects hundreds of lower-affinity interactions missed by PBM and HT-SELEX. **(A)** The PADIT-seq genetic circuit is shown with a reporter library containing all possible 10-bp DNA sequences as candidate TFBS. **(B)** The PADIT-seq reporter library was mixed with either a ‘no DBD’ control or a constitutive T7 promoter driving expression of either H0XD13 or EGR1. **(C)** Scatter plot of counts per TFBS between 2 replicates of PADIT-seq experiments with HOXD13 (left) and EΘR1 (right). **(D)** EGR1 PA-DIT-seq activities are compared to MITO-Ml-derived dissociation constants (K_d_). **(E)** PADIT-seq activities are compared to PBM E-scores for all possible 8-mers for HOXD13 (left) and 9-mers for EGR1 (right). Active TFBS are red, inactive TFBS are black. **(F)** Plots in panel D are shown again with blue indicating TFBS that are significantly enriched in HT-SELEX cycle 4 for H0XD13 and cycle 2 for EGR1, which were determined to be appropriate for PWM generation in Jolma et al. Cell, 2013. **(inset)** PADIT-seq activities for active TFBS enriched (blue) or not enriched (red) in HT-SELEX. * denotes a Wilcoxon rank sum test p-valυe < 0 05

### PADIT-seq detects hundreds of lower affinity interactions missed by PBM and HT-SELEX

To benchmark PADIT-seq, we assayed two extensively studied and well-characterized human TFs: EGR1 and HOXD13, representing the largest (Cys_2_His_2_ (C2H2)-type zinc finger) and second largest (homeodomain) TF families, respectively, in mammals. EGR1 is an immediate early gene and a major regulator of synaptic plasticity and neuronal activity (*40*). HOXD13 is a critical regulator of limb development, and mutations in HOXD13 have been linked with various limb malformations (*41*). To assay these TFs by PADIT-seq, we first constructed a reporter library with all possible 10-bp DNA sequences as candidate TFBS (n = 1,048,576) (Methods). TFBS were randomly associated with BCs during PADIT-seq reporter library construction, and TFBS-BC combinations were determined by Illumina DNA sequencing. The PADIT-seq reporter library was then mixed with either a ‘no DBD’ control or a constitutive T7 promoter driving expression of either HOXD13 or EGR1 (Figure 1B). Following *in vitro* transcription and translation (IVTT), Illumina sequencing of reporter RNAs was performed, and BC counts per TFBS for HOXD13, EGR1 and the ‘no DBD’ control were found to be highly reproducible across triplicate assays (Figure 1B, Figure S1).

To quantify the TF-DNA interactions, we performed differential gene expression analysis of TFBS counts against the ‘no DBD’ control using DESeq2 (*42*); we defined the resulting log_2_ (DBD / ‘no-DBD’) values from DESeq2 as the TF DBD’s ‘PADIT-seq activity’. We defined ‘active’ TFBS as those that significantly increased reporter gene expression upon TF binding. At a false discovery rate (FDR) of 5%, we identified 46,279 and 6,596 active 10-mers for HOXD13 and EGR1, respectively. To test the robustness of PADIT-seq in identifying active *k*-mers, we constructed a smaller-scale PADIT-seq reporter library comprising 896 candidate 9-bp binding sites for HOXD13 or EGR1 (Methods). For both TFs, all of the PADIT-seq active *k*-mers identified in the all 10-mers library were also active in this smaller-scale library (Figure S2 A-B). PADIT-seq activities were also highly correlated between the all 10-mers and smaller-scale library for both HOXD13 (Figure S2A; Pearson r = 0.959) and EGR1 (Figure S2B; Pearson r = 0.974), supporting the reliability of PADIT-seq results.

To assess the accuracy of PADIT-seq in quantifying TF-DNA interaction strengths, we compared EGR1 PADIT-seq activities to equilibrium dissociation constants (K_d_) derived an orthogonal MITOMI assay (*43*). PADIT-seq activities for EGR1 from both libraries were highly correlated with MITOMI-derived K_d_ values (Figure 1C, S2C). Importantly, EGR1 binding to low affinity sites (K_d_ ∼ 0.1 μM) was detected among the active *k*-mers in both libraries (Figure 1C, S2C).

To further assess the accuracy of PADIT-seq in identifying genuine low affinity DNA binding sites, we performed an in-depth comparison of PADIT-seq activities from the all-10mers library to PBM enrichment (E) scores. Since PBM provides a TF binding score to all possible 8-mers and 9-mers, we first derived median 8-mer and 9-mer PADIT-seq activities for HOXD13 and EGR1, respectively, from the all-10mers data (Methods). HOXD13 was active at 1,780 of the 32,896 possible 8-mers (∼5.4%). In contrast, EGR1, which recognizes DNA sequences approximately 9-bp in length (*44*), was active at 554 of the 131,072 possible 9-mers (0.42%). PBM and PADIT-seq results were highly concordant; HOXD13 and EGR1 active sites represented the top 1,780 and 554 PBM 8-mers and 9-mers, respectively. However, the PBM E-score cutoffs for differentiating these top bound sites were quite different for the two TFs (0.30 for HOXD13 and 0.45 for EGR1; Figure 1D), suggesting that using a fixed PBM threshold across TFs does not reliably discriminate bound sites.

We next evaluated the ability of HT-SELEX, which like PBMs (*3, 5*), has been used to assay the DNA binding preferences of hundreds of TFs (*45*), for its ability to distinguish PADIT-seq active *k*-mers. We re-analyzed publicly available HOXD13 and EGR1 HT-SELEX data (*45*) (Methods) and identified 196 8-mers and 83 9-mers enriched for HOXD13 and EGR1 binding, respectively, which are >5-fold fewer active *k*-mers identified by PADIT-seq. For both TFs, the HT-SELEX enriched *k*-mers were biased for detecting high affinity TFBS and missed lower affinity interactions identified by PADIT-seq (Figure 1E), irrespective of the HT-SELEX cycle analyzed (Figure S3). PWMs derived from HT-SELEX data were similarly biased for detecting high affinity interactions (Figure S4 and S5) (*46, 47*). Therefore, PADIT-seq significantly outperforms existing high-throughput technologies in identification of lower affinity TF-DNA binding sites.

### Nucleotides flanking high affinity binding sites create overlapping lower affinity sites

To investigate the role of lower affinity binding sites in TF occupancy *in vivo*, we next used PADIT-seq data to inspect genomic regions bound by Hoxd13 in the murine embryonic forelimb bud (*48*) and by Egr1 in the murine frontal cortex (*49*). The DBD amino acid sequences of HOXD13 and EGR1 are identical between mouse and human. We first asked whether incorporating lower affinity TF-DNA interactions increases the ability to discriminate ChIP-seq ‘bound’ regions from background genomic regions. For Egr1, both PADIT-Seq *k*-mer data and a PWM derived from HT-SELEX data performed very well (AUROC ∼0.97) (Figure S6A right), indicating that high affinity binding sites are sufficiently predictive of *in vivo* Egr1 binding. In contrast, for Hoxd13, both PADIT-Seq *k*-mer data and a PWM derived from HT-SELEX data performed only moderately well (AUROC ∼0.77) (Figure S6A left), consistent with the known role of protein cofactors in modulating Hox factor binding *in vivo* (*7, 50*).

Although lower affinity sites did not aid in discriminating sites bound by Egr1 or Hoxd13 *in vivo*, we hypothesized that they might have a quantitative effect on the level of TF binding *in vivo*. For both Egr1 and Hoxd13, we found that integration of all the PADIT-seq active *k*-mers’ activity levels within the ChIP-seq peaks was significantly correlated with normalized ChIP-Seq read counts (Pearson r ∼0.32; *P* < 2.2 x 10^-16^ for both TFs; Figure S6B). Considering only the top active *k*-mers, corresponding to the highest affinity TFBS, resulted in lower correlation with ChIP-seq signal than when lower affinity TFBS were included in the analysis (Figure S6C-D).

To investigate how lower affinity binding sites increase TF genomic occupancy, we performed a sliding window analysis in which we scored every 8-mer (for Hoxd13) or 9-mer (for Egr1), moving in 1-bp steps, across Hoxd13 or Egr1 ChIP-seq peaks, respectively. We found multiple, consecutive active *k*-mers within both Hoxd13 and Egr1 ChIP-seq peaks. For example, at the ChIP-seq peaks near *Cadps* and *Lncbate10*, there were 6 consecutive, active Hoxd13 8-mers and 4 consecutive, active Egr1 9-mers, respectively (Figure 2A). This phenomenon, where multiple TFBS overlap each other, is distinct from homotypic clustering, in which multiple, non-overlapping binding sites for a particular TF are located typically tens to hundreds of bp apart, within a *cis*-regulatory element (*51–53*). We investigated further because to our knowledge, this phenomenon has not been reported previously.

**Figure 2:**
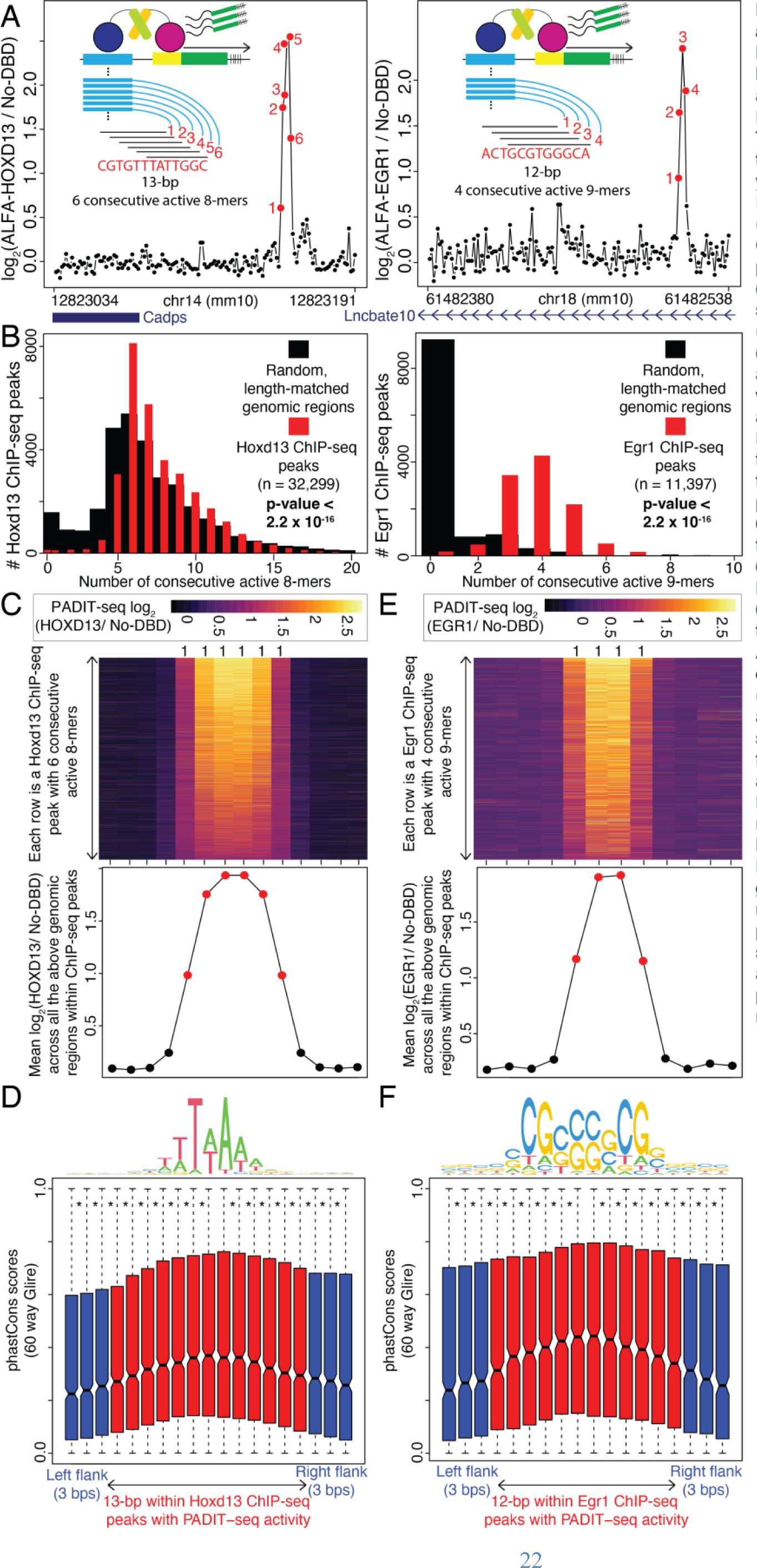
Nucleotides flanking high affinity TFBS create overlapping lower affinity binding sites. **(A)**k-mers are tiled in 1 -bp steps across an example Hoxd13 {left; 8-mers) and Egr1 (right; 9-mers) ChIP-seq peak. The corresponding PADIT-seq activi-ties for every k-mer is plotted on the y-axis. Active k-mers are colored red; Inactive k-mers are colored black, **(inset)** Every k-mer tiled across ChIP-seq peaks was profiled inde-pendently In the PADIT-seq assay. **(B)** Histogram of the number of con-secutive active k-mers in Hoxd13 (left) and Egr1 (right) ChIP-seq peaks. **(C)** (Top) Heatmap of PADIT-seq activity at Hoxd13 ChIP-seq peaks with 6 consecutive active 8-mers, along with 4 flanking inactive 8-mers Each row is a ChIP-seq peak. (Bot-tom) Mean PADIT-seq activities of all the genomic regions within ChIP-seq peaks in the heatmap above. **(D)** 60-way Glire PhastCons scores for the 13-bp genomic regions containing 6 consecutive active 8-mers in Hoxd13 ChIP-seq peaks (red). 60-way Glire PhastCons scores for the flanking 3 nucleotides are in blue. Adjusted p-values from paired Wilcox-on tesls < 0.05 are indicated by ‘. **(E)** (Top) Heatmap of PADIT-seq activity at Egr1 ChIP-seq peaks with 4 con-secutive active 9-mers, along with 4 flanking ‘inactive’ 9-mers. Each row is a ChIP-seq peak. (Bottom) Mean PA-DIT-seq activities of all the genomic regions within ChIP-seq peaks in the heatmap above. **(F)** 60-way Glire PhastCons scores for the 12-bρ genomic regions containing 4 consec-utive active 9-mers in Egrī ChIP-seq peaks (red). 60-way Glire PhastCons scores for the tlanking 3 nucleotides are in blue. Adjusted p-values from paired Wilcoxon tests < 0.05 are marked by *.

Across all Hoxd13 and Egr1 ChIP-seq peaks, we found that Hoxd13 and Egr1 ChIP-seq peaks were significantly enriched for having a larger number of consecutive, active *k*-mers (Wilcoxon rank sum test *P* < 2.2 x 10^-16^ for both TFs; Figure 2B and S7). ∼87% of Hoxd13 ChIP-seq peaks contained >5 consecutive, active 8-mers, and ∼63% of Egr1 ChIP-seq peaks contained >3 consecutive, active 9-mers, both of which are significantly more than in background genomic regions (Figure 2B and S7). Across all the Hoxd13 ChIP-seq peaks with ≥6 consecutive active 8-mers, a high affinity active 8-mer tended to be the central TFBS and was flanked by nucleotides that created overlapping, lower affinity Hoxd13 binding sites (Figure 2C and S8). To further investigate whether the consecutive, active 8-mers may be functionally important, we inspected the evolutionary conservation of the corresponding genomic regions. We found that the 13-bp genomic regions corresponding to 6 consecutive, active Hoxd13 8-mers (n = 8,122 ChIP-seq peaks) were significantly more conserved than the flanking genomic regions (Figure 2D; paired Wilcoxon rank sum test adjusted *P* < 0.05); this higher level of conservation was robust to variation in the number of consecutive, active 8-mers (Figure S8). Similarly, among Egr1 ChIP-seq peaks (n = 11,397), 4 consecutive active 9-mers were observed most frequently (n = 4,271; ∼37%). 9-mers with high affinity for Egr1 were flanked by nucleotides that created additional, overlapping lower affinity 9-mer binding sites (Figure 2E and S8), and the corresponding genomic regions were more conserved than the flanking nucleotides (Figure 2F and S9).

Since some TFs prefer binding sites with particular DNA shape features (*e.g.*, narrow minor groove width (*54*)), we investigated whether the consecutive active *k*-mers found within Hoxd13 and Egr1 ChIP-seq peaks are enriched for any DNA shape features. Using a deep learning based tool, Deep DNAShape (*55*), we found that minor groove width (Figure S10) and propeller twist (Figure S11) at the extended recognition sequences bound by Hoxd13 and Egr1 were distinct from flanking genomic regions. This is consistent with a previous study where distinct DNA shapes at nucleotides flanking motif matches influenced TF occupancy (*14*). However, given that Hoxd13 and Egr1 binding to AT- and GC-rich binding sites, respectively, which are known to have distinct DNA shapes (*54*), further studies are required to better understand the relationship between TF DNA shape preferences and lower affinity binding sites uncovered here.

### Overlapping binding sites increase TF occupancy *in vitro*

We performed three analyses to test the model that we derived from analysis of ChIP-Seq and PADIT-Seq data, *i.e.*, that consecutive, active *k*-mers, overlapping by 1-bp, increase TF occupancy. First, we analyzed HT-SELEX data. Since every round of selection in HT-SELEX enriches DNA sequences with higher affinity for the assayed TF, our model predicts that the number of consecutive, active *k* -mers should increase in frequency after every round of selection, which is indeed what we observed for both HOXD13 (Figure 3A) and EGR1 (Figure 3B).

**Figure 3:**
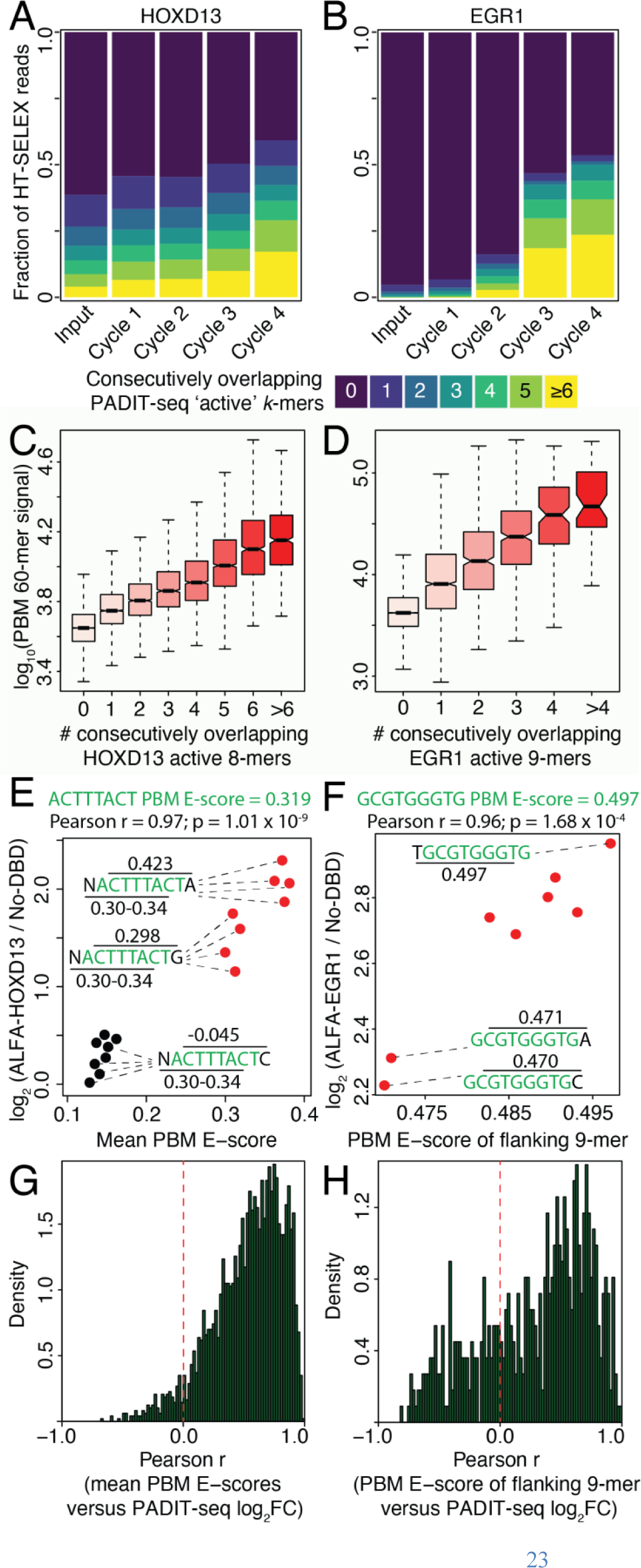
Overlapping binding sites increase TF occupancy *in vitro*. **(A-B)** The fraction of HT-SELEX reads (y-ax-is) with consecutive overlapping active 8-mers and 9-mers for HOXD13 (A) and EGR1 (B), respectively, after 0-4 rounds of selection (x-axis). **(C-D)** 60-bp PBM probes (n = −42Ό00) are categorized by the number of consecutive overlapping PADIT-seq active 8-mers and 9-mers for HOXD13 (C) and EGR1 (D), respec-tively. The corresponding signal intensi-ties are plotted on the y-axis. Wilcoxon tests for all pairwise comparisons have an adjusted p-value < 0.05 (not indicat-ed). **(E)** 10-mer HOXD13 PADIT-seq activiy of all possible 1-bp nucleotides flanking the 8-mer ‘ACTTTACT’, against the mean PBM E-scores of the 3 con-secutive 8-mers comprising each 10-mer. **(F)** 10-mer EGR1 PADIT-seq activity of all possible 1-bp nucleotides flanking the 9-mer ‘GCGTGGGCG’, against the PBM E-score of the remain-ing 9-mer comprising each 10-mer. **(G)** For all active 8-mers, histogram of the Pearson correlation coefficients between 10-mer HOXD13 PADIT-seq activity and the mean PBM E-scores of the 3 consecutive 8-mers comprising each 10-mer. **(H)** For all active 9-mers, histogram of the Pearson correlation coefficients between 10-mer EGR1 PA-DIT-seq activity and the PBM E-score of the remaining 9-mer comprising each 10-mer.

Second, our model predicts that PBM signal intensities should increase with increasing numbers of consecutive, active *k*-mers. When PADIT-seq *k*-mers were tiled in 1-bp steps across ∼42,000 60-bp PBM probes assayed for binding by HOXD13 or EGR1, we found that the PBM signal intensities for the 60-mer probes were directly proportional to the number of consecutive, active 8-mers and 9-mers for both HOXD13 (Figure 3C) and EGR1 (Figure 3D), respectively. Overlapping binding sites had a significantly higher effect on PBM signal intensities than equivalent numbers of non-overlapping binding sites (Figure S12).

Third, our model predicts that HOXD13 10-mer PADIT-seq data should be better correlated to the sum of PBM E-scores of all three 8-mers overlapping each 10-mer, instead of any single 8-mer PBM E-score. For example, for all 10-mers with a central ATTTTATT 8-mer, the mean PBM E-scores of the 2 overlapping 8-mers were highly correlated with the 10-mer PADIT-seq activities (Figure 3E). Similarly, while the 9-mer GCGTGGGTG is a high-affinity binding site for EGR1 (MITOMI K_d_ = 17.4 nM (*43*); PBM E-score = 0.497), nucleotides flanking it create an additional 9-mer whose PBM E-scores (0.47 < E < 0.498) are highly correlated with the corresponding 10-mer PADIT-seq activities (Figure 3F). A positive correlation between mean PBM E-scores of overlapping *k*-mers and 10-mer PADIT-seq activities was found for the majority of active *k*-mers for both TFs (Figure 3G-H; binomial test *P* < 2.2 x 10^-16^).

Altogether, we observed strong quantitative agreement between all 3 predictions of our model and empirical measurements, providing strong support for the additive roles of consecutive active *k*-mers in modulating TF-DNA binding.

### ‘Weavability’ of active TF binding sites

Our results on overlapping TFBS led us to ask: how are consecutive, active *k*-mers interwoven to create extended recognition sites? To answer this question, we constructed a network in which nodes correspond to HOXD13 active 8-mers and edges between nodes indicate a 7-bp overlap between the connected 8-mers (Figure 4A). We found that 97% of HOXD13 active 8-mers formed a single connected network (Figure 4B), indicating an inherent capacity of HOXD13 binding sites to weave together into extended recognition sequences. Similarly, 88% of EGR1 active 9-mers formed a single connected network (Figure 4C). Moreover, for both TFs, we found that the binding affinity of a node (*i.e.*, the *k*-mer’s PADIT-seq activity) was positively correlated with the total number of its edges (Figure 4D and Figure 4E). These results suggest that high affinity *k*-mers are more likely to be flanked by nucleotides that create overlapping binding sites than lower affinity *k*-mers, explaining why high affinity active *k*-mers tend to be the central TFBS in Hoxd13 and Egr1 ChIP bound regions (Figure 2C and 2E).

**Figure 4:**
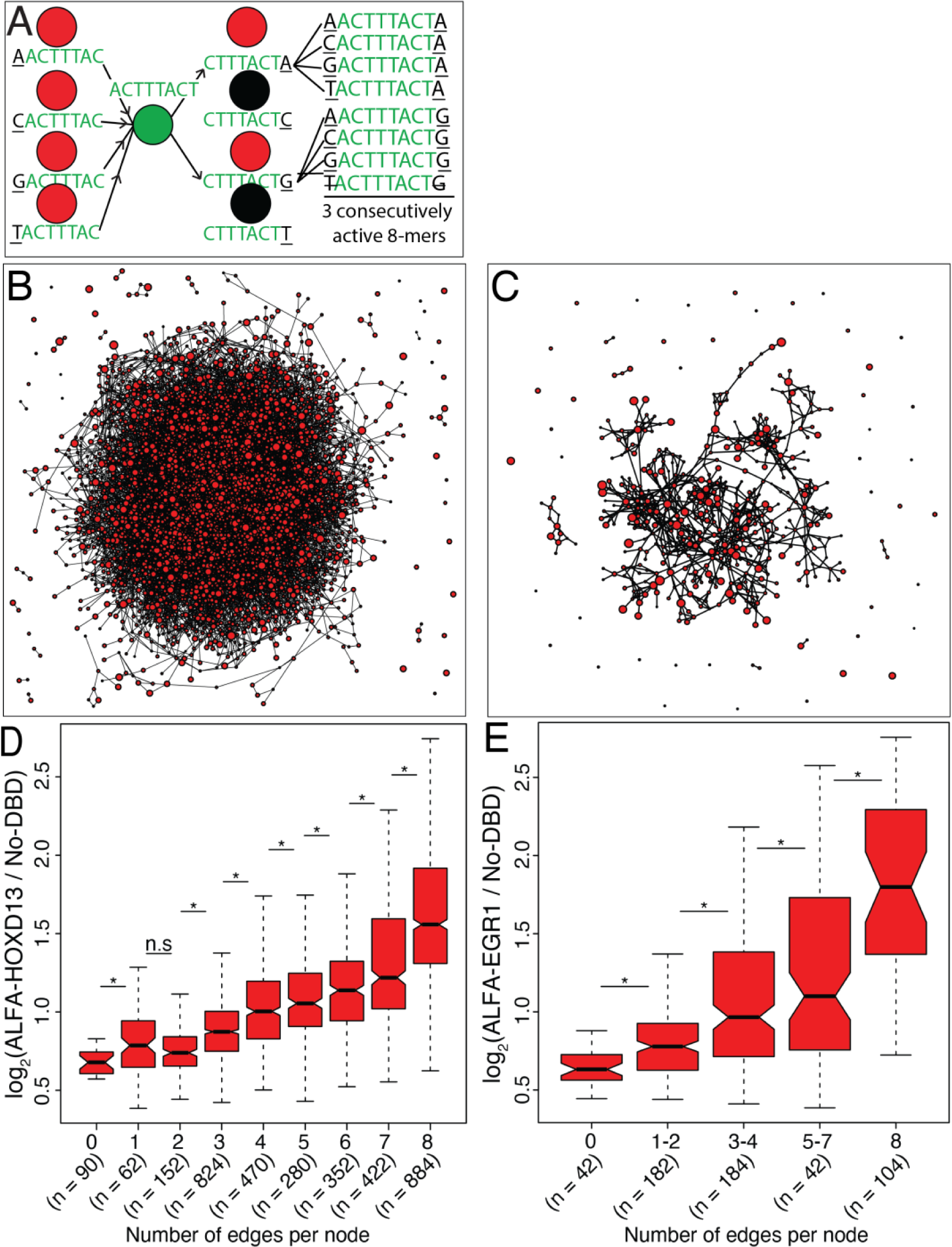
Active HOXD13 and EGR1 k-mers are inherently ‘weavable’. **(A)** Schema to demonstrate the logic of network construction in panel B Active 8-mers are colored red, inactive 8-mers are colored black. **(B)** Network representation of H0XD13 active 8-mers (n = 3,536) connected by directed edges (arrows not shown). The logic of network construction is demonstrated in panel b. 3,446 out of 3,536 nodes (97.4%) form the largest, single connected component. **(C)** Network representation of EGR1I active 9-mers (n = 554) connected by directed edges (arrows not shown). 487 out of 554 nodes (∼87.9%) form the largest, single connected component. **(D-E)** PADIT-seq activity of active 8-mers (HOXD13, panel C) and 9-mers (EGR1, panel E) is plotted against the total number of incoming and outgoing edges per node. Adjusted p-values from Wilcoxon tests < 0.05 are marked by *.

**Figure 5:**
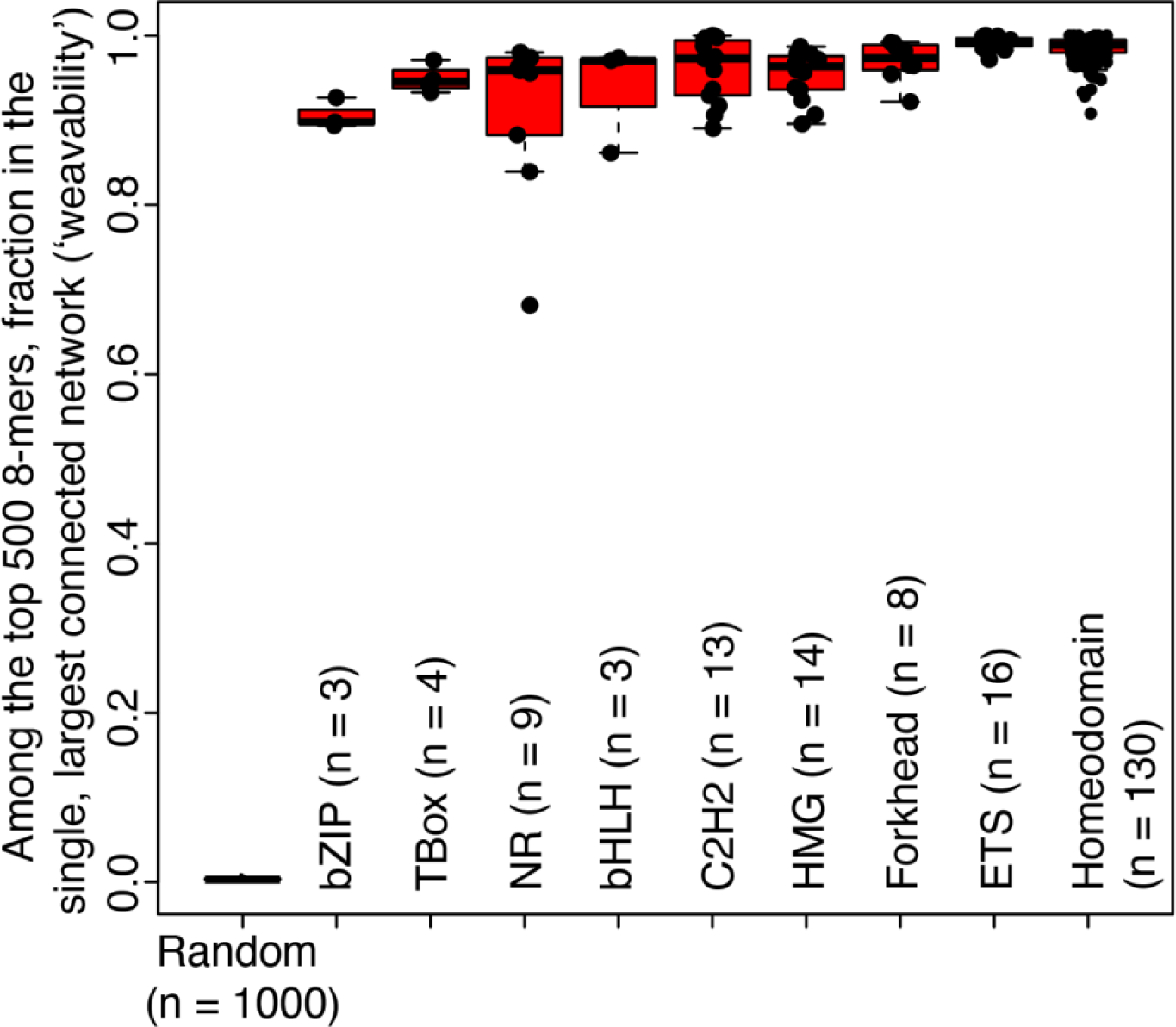
‘Weavability’ of binding sites is an inherent property of TFs from different DBD classes. Among the top 500 PBM 8-mers, including reverse complements, the fraction of nodes in the single, largest connected network is plotted for 200 TFs from 9 different families of DBDs. 1000 random samplings of 500 8-mers are also shown (‘Random’), and were used to perform the Permutation Test.

Next, we investigated whether the ‘weavability’ of TF binding sites is a more general phenomenon, beyond HOXD13 and EGR1. In the absence of PADIT-seq data for additional TFs, we leveraged the PBM datasets publicly available in the UniPROBE database for hundreds of TFs from diverse families of DBDs (*56, 57*). Even though fixed PBM E-score thresholds do not reliably differentiate bound versus unbound sites (Figure 1D), PADIT-seq active *k*-mers were highly concordant with the corresponding top PBM *k*-mers for both HOXD13 and EGR1. Therefore, we constructed directed networks for the top 500 8-mers with the highest PBM E-scores for 200 different human and mouse TFs from 9 different TF DBD classes (bZIP, T-Box, Nuclear Receptors, bHLH, C2H2-type zinc fingers, HMG, Forkhead, ETS, Homeodomain). The majority of the top 500 8-mers formed a single, highly connected network for all 200 TFs (permutation test *P* < 0.001 for each TF), suggesting that the ‘weavability’ of TF binding sites is an intrinsic property of diverse metazoan TF classes.

### Noncoding variants alter multiple overlapping binding sites to influence TF binding and gene expression

We investigated the impact of noncoding variants on overlapping TF binding sites and gene expression *in vivo*. We analyzed the allelic skew of reporter gene expression in massively parallel reporter assays (MPRA) for variants predicted by PADIT-seq to alter EGR1 binding. We focused this analysis on EGR1 because HOXD13 has restricted gene expression in adult cells (*58*), while EGR1 is widely expressed across many different cell types, with important roles in differentiation (*59*), memory formation (*60*), insulin secretion (*61*) and immune response (*62*).

We analyzed 4,784 variants for which allelic skew was detected in four published MPRAs performed in neurons (*63*), pancreatic beta cells (*64*) and lymphoblastoid cell lines (*65, 66*). Of these, we observed differential PADIT-seq activity at 149 variants (Methods). Alleles with higher PADIT-seq activity showed higher MPRA activity for 91 variants and lower MPRA activity for 58 variants (Figure 6A; binomial test *P* = 0.00853), consistent with EGR1’s known roles as both a transcriptional activator and repressor (*67, 68*).

**Figure 6:**
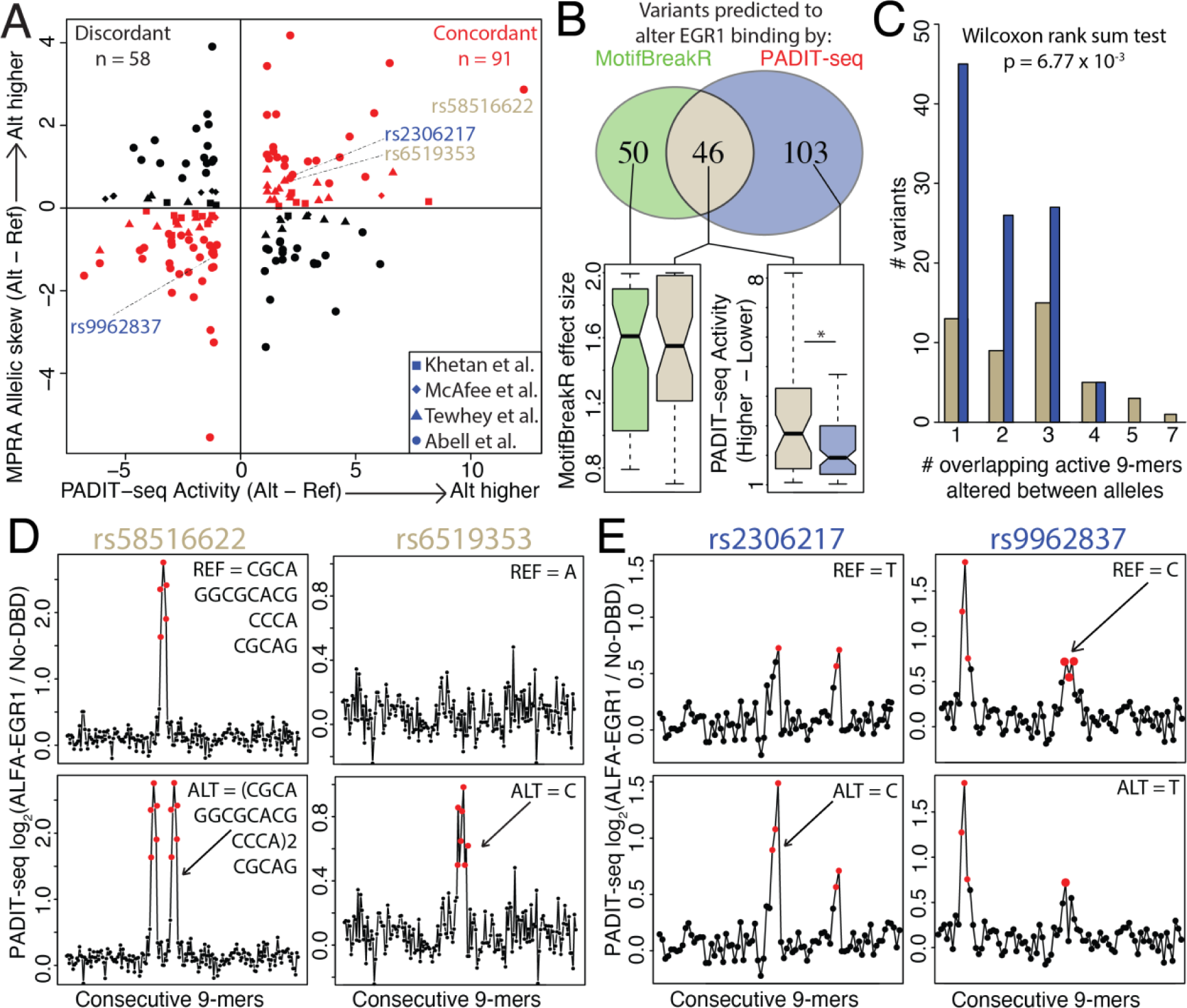
Noncoding variants alter multiple overlapping EGR1 binding sites to influence gene expression. **(A)** Allelic skew in MPRA activity of 149 variants with differential EGR1 PADIT-seq activity. Shapes represent the different studies from which allelic skew in MPRA activity was obtained. Red shapes correspond to concordant directions of effect between differential EGR1 PADIT-seq activity and MPRA allel-ic skew. **(B)** Euler diagram of variants with MPRA allelic skew predicted by MotifBreakR to alter EGR1 bind-ing (green) and with differential PADIT-seq activity (blue). (Bottom) Boxplots comparing MotifBreakR (left) and PADIT-seq (right) effect sizes. * indicates Wilcoxon rank sum test p-value < 0.05. **(C)** Number of active overlapping 9-mers altered by the 149 variants with differential PADIT-seq activity. Brown variants are pre-dicted to alter EGR1 binding by MotifBreakR, while blue are not. **(D-E)** Reference (Top) and alternate (Bot-tom) alleles of 4 variants, highlighted in panel A, are tiled in 1-bp steps, and the corresponding EGR1 PA-DIT-seq activities for every 9-mer are plotted on the y-axis. All 4 variants have differential PADIT-seq activi-ty, while only the variants in panel D are predicted by MotifBreakR to alter EGR1 binding.

We next compared differential PADIT-seq activity to a popular PWM based algorithm that scores the effects of variants on TF binding, MotifBreakR (*69*). While ∼31% of variants with differential PADIT-seq activity were also predicted to alter EGR1 binding by MotifBreakR, the majority of them were missed (Figure 6B). These variants had lower effects on EGR1 binding (Wilcoxon rank sum test *P* = 0.033; Figure 6B) and altered fewer overlapping binding sites (Wilcoxon rank sum test *P* = 6.77 x 10^-3^; Figure 6C). For example, variants rs58516622 and rs6519353 (Figure 6D), which are predicted by MotifBreakR to alter EGR1 binding, have large effects because they alter 5 and 7 overlapping binding sites, respectively. In contrast, variants, rs2306217 and rs9962837, alter 2 overlapping binding sites each and are missed by MotifBreakR. Our results, therefore, suggest that noncoding variants influence gene expression by modulating multiple, consecutive active *k*-mers.

To directly investigate how noncoding variants modulate consecutive active *k*-mers, we analyzed 5,748 and 4,136 variants that were directly tested by SNP-SELEX for differential binding to HOXD13 and EGR1, respectively (*70*). Using PADIT-seq data, we accurately identified 92.8% (39/42) and 96.4% (81/84) of variants found by SNP-SELEX to alter HOXD13 and EGR1 binding, respectively (Figure 7A). These variants primarily had large effects on TF binding (brown, Figure 7A). However, PADIT-seq data indicated that many more variants would cause differential binding (Figure 7A insets): 1,880/5,748 for HOXD13 and 761/4,136 for EGR1. Although these variants were not identified as differentially bound by SNP-SELEX (yellow, Figure 7A), the effects of these variants on PADIT-Seq activity were highly correlated with SNP-SELEX preferential binding scores (Figure 7A), suggesting they are likely true positives, but below the sensitivity of SNP-SELEX to detect them. Notably, the SNP-SELEX preferential binding scores for all assayed variants were highly correlated with the number of consecutive, active *k*-mers that were altered between alleles (Figure 7B), consistent with the model that noncoding variants alter TF binding by perturbing multiple, overlapping binding sites.

**Figure 7:**
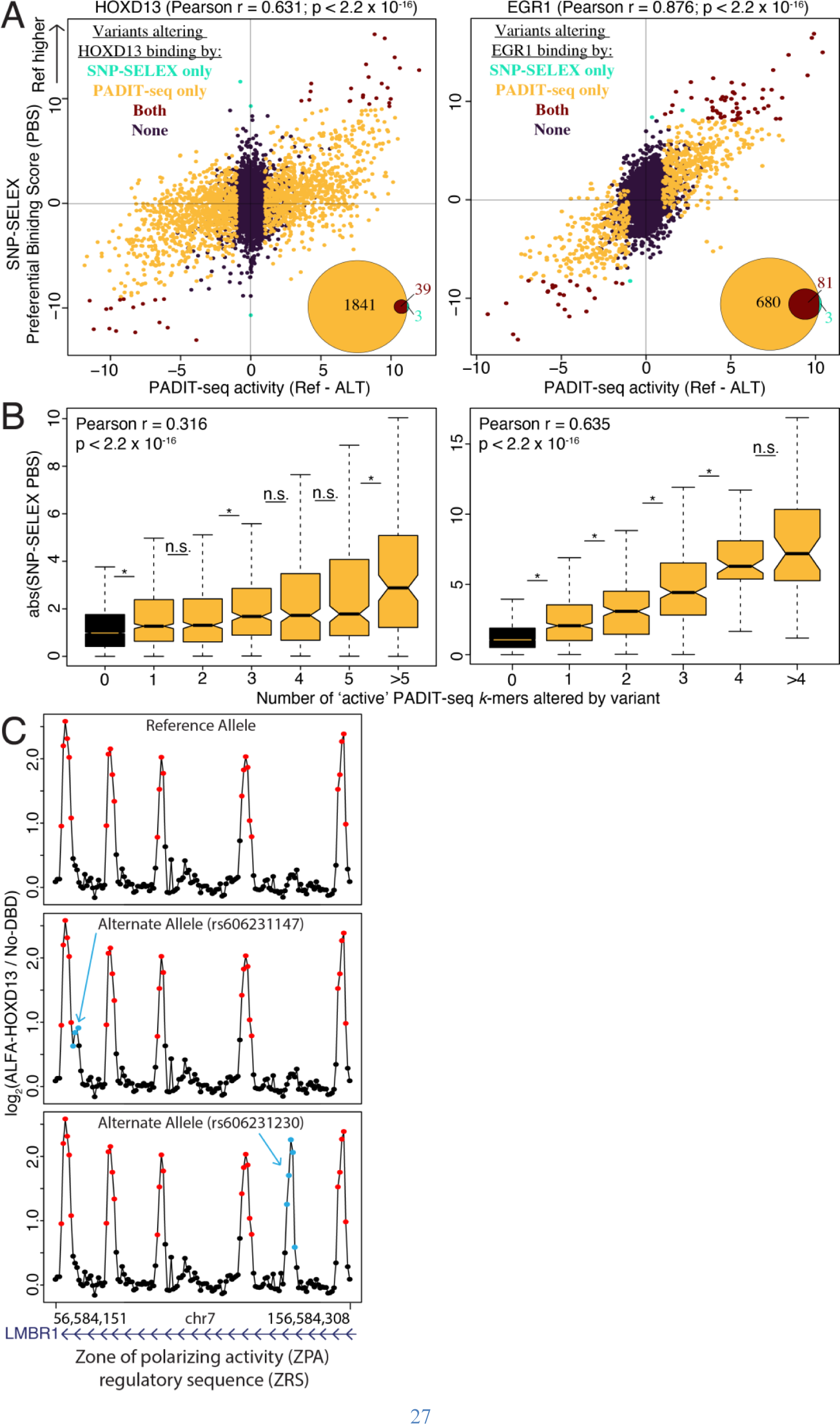
Noncoding variants alter TF binding by perturbing multiple overlapping binding sites. **(A)** Difference in PADIT-seq activity between k-mers tiled across the reference and alternate alleles (x-axis) is plotted against the SNP-SELEX preferential binding scores (y-axis) for H0XD13 (left) and EGR1 (right). Pearson correlation coefficients for all variants except those colored black are indicated, **(inset)** Euler dia-gram of variants with differential PADIT-seq activity and SNP-SELEX preferential binding scores. **(B)** Number of overlapping PADIT-seq active k-mers altered by variant alleles is plotted against the SNP-SELEX preferential binding scores. **(C)** Human genetic sequence corresponding to the zone of polar-izing activity (ZPA) regulatory sequence (ZRS), which is a limb-specific enhancer occupied by Hoxd13 in the developing mouse limb bud, is tiled in 1 -bp steps, and HOXD13 PADIT-seq activities are plotted on the y-axis for the reference allele (Top), and risk alleles (Bottom) for preaxial polydactyly. Active 8-mers are colored red; risk allele specific active 8-mers are col-ored blue; Inactive 8-mers are colored black.

Given the utility of PADIT-seq data in identifying variants that alter TF binding, we investigated whether noncoding variants associated with various limb malformations might alter HOXD13 DNA binding. Human coding mutations in HOXD13 have been implicated in syndactyly (*71*), brachydactyly (*72*) and synpolydactyly (*73*), and 13 different noncoding variants, annotated as pathogenic for various limb malformations in the ClinVar database (*74*), overlap a limb-specific enhancer (*75*) occupied by Hoxd13 in the developing mouse limb bud (*48*). A sliding window analysis identified 2 putatively causal variants for preaxial polydactyly, rs606231147 and rs606231230, that have higher PADIT-seq activity for the risk allele by creating multiple, overlapping HOXD13 active 8-mers (Figure 7C). This is consistent with a recent study, which found that other noncoding variants in this limb-specific enhancer, putatively causal for polydactyly, increased binding by ETS factors (*76*). Our results, therefore, suggest that alterations to multiple, consecutive binding sites might be a general mechanism by which noncoding variants affect gene regulation and human traits, including diseases.

## Discussion

PADIT-seq overcomes limitations of prior high-throughput approaches for assaying TF-DNA interactions by enabling sensitive detection of both high and lower affinity DNA binding sites. The expanded repertoire of bound sequences uncovered by PADIT-seq provides key insights into the role of lower affinity sites in modulating TF occupancy *in vivo*. We found that lower affinity binding sites tend to flank high affinity *k*-mers, creating extended recognition sequences in most ChIP bound genomic regions. Recent independent work showed that partition function models that sum binding contributions from short tandem repeats (STRs) flanking core motifs are predictive of overall TF binding (*33*). However, a key difference between those results and the model that we present in this study is that while multiple TF molecules can bind a given DNA sequence when STRs are present on that DNA fragment, we focused on the effects of overlapping binding sites on binding by a single TF molecule. Moreover, the extended recognition sequences are not limited to STRs, which overlap only ∼8% of the Hoxd13 and Egr1 ChIP-seq peaks but are present in almost all of the bound genomic regions.

We propose a model in which nucleotides adjacent to high affinity sites create overlapping, lower affinity sites that together influence TF binding (Figure 8). Formation of such extended recognition sequences stems from an inherent property of TF binding sites to interweave each other. Implicit in the ‘weavability’ of TF binding sites is the interdependence of nucleotide positions within TF binding sites. Moreover, PADIT-seq has revealed a much wider range of lower affinity binding sites to consider when interpreting the functional consequences of noncoding variants, which we show modulate affinity through the cumulative effects on multiple, overlapping binding sites.

**Figure 8:**
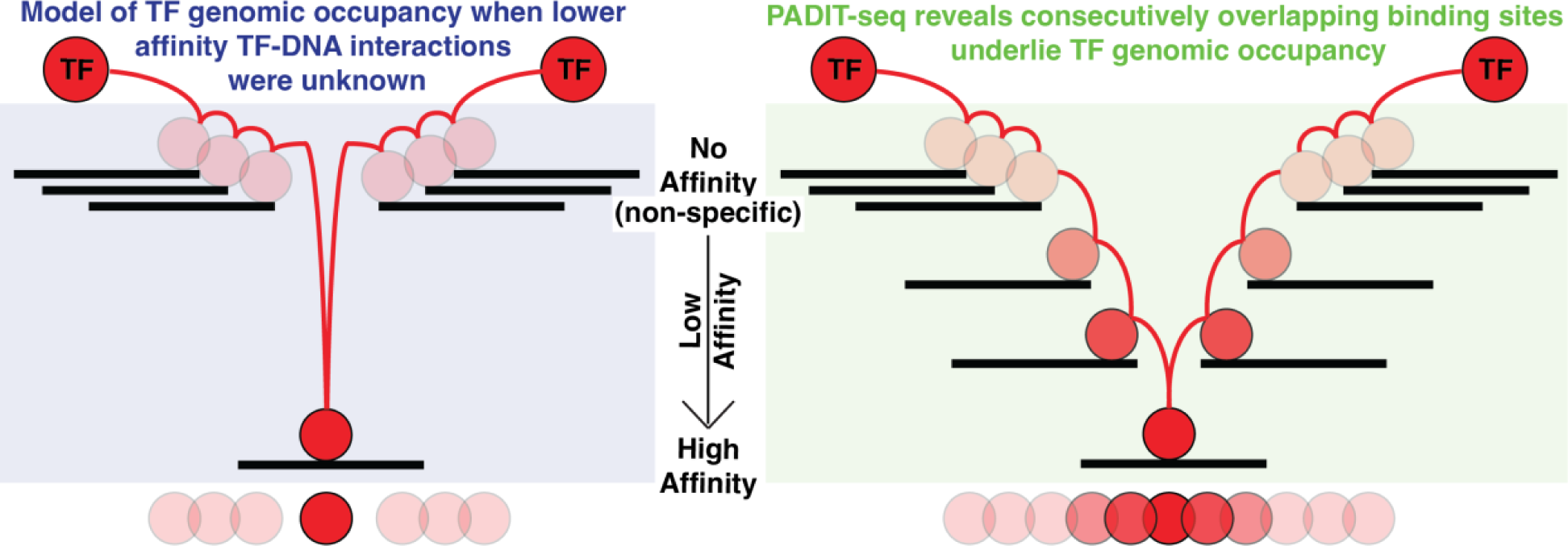
PADIT-seq reveals consecutively overlapping binding sites underlie TF genomic occupan-cy. The significantly expanded repertoire of lower affinity binding sites uncovered by PADIT-seq reveals that high affinity sites are flanked by nucleotides that create additional, overlapping lower affinity binding sites that together influence genomic TF occupancy.

Altogether, our results revealed unanticipated complexity in the rules governing TF-DNA interactions and suggest that the intrinsic property of high affinity sites to have overlapping, lower affinity sites may confer them with a previously unappreciated mechanism by which they provide an energetic ‘sink’ for TF binding. Future studies are needed to determine how the intrinsic weavability of TF binding sites may have evolved. Overlapping DNA binding sites may enable a more nuanced control of TF binding and target gene expression. As machine learning methods are increasingly being developed to decipher *cis*-regulatory codes (*77–79*) and design synthetic *cis*-regulatory elements (*80*), consideration of consecutive overlapping binding sites may improve model performance.

### Limitation of PADIT-seq

To mitigate potentially confounding impacts when interrogating all possible 10-bp DNA sequences as candidate TFBS, the flanking nucleotides were intentionally selected to preclude all adjoining 8-mers with HOXD13 and EGR1 PBM E-scores >0.25. This stringent E-score cutoff of 0.25 for flanking 8-mers proved prudent given the detected influence of adjacent lower affinity sites. However, this also means that surveying binding landscapes for new TFs using PADIT-seq necessitates consideration of potential impacts from sequences flanking candidate TFBS.

## Acknowledgments

We thank Steve Gisselbrecht for help with calculating PBM 9-mer E-scores for EGR1. We thank members of the Bulyk lab for helpful discussions, and Luca Mariani, Katrina Liu, and Brent Carroll for critically reading the manuscript. This work was supported by the National Institutes of Health (grants R21 HG010200 and R01 HG010501 to M.L.B.).

## Author contributions

M.L.B. conceived the research project. M.L.B. and S.K. designed the research project. S.K. performed experiments and analyses and prepared the figures. M.L.B. supervised the research. M.L.B. and S.K. wrote the manuscript. All authors approved the final version of the manuscript.

## Data and materials availability

PADIT-Seq data have been deposited in the GEO database under accession number GSE250601. Code and processed data for generating the figures are available at https://github.com/BulykLab/PADIT-seq.

## Materials and Methods

### Cloning DBDs into the pET28 vector

The low-copy pET28 bacterial expression vector was used as a template for two high-fidelity Q5 polymerase PCR reactions. Primers ‘pET28_pcr2_FWD’ and ‘pT7LACI_REV’ were used to generate a 2823-bp fragment, and primers ‘T7_Terminator_FWD’ and ‘pET28_pcr1_REV’ were used to generate a 2260-bp fragment (see Table S1 and S2). The 2 pET28 amplicons along with the ALFA-DBD DNA sequence, obtained as an IDT gBlock gene fragment, were combined to construct the plasmid using 3-fragment Gibson assembly. The reaction mixture was transformed into TOP10 *E. coli*. Multiple colonies were Sanger sequenced to identify clones with correct assembly and sequence. The region encoding the ALFA-DBD fusion, along with the T7 promoter and Terminator, was PCR amplified using Q5 polymerase and primers ‘T7_Promoter_Fwd’ and ‘T7_Terminator_REV’. This ‘pT7-ALFA-DBD-T7Term’ amplicon was purified with QiaQuick PCR purification kit and used as a template in an *E. coli*-based PURExpress *in vitro* transcription and translation (IVTT) reaction. Expression of the ALFA-DBD protein was validated by SDS-PAGE and western blotting using an anti-FLAG antibody.

### Cloning nbALFA-T7-RNA-Polymerase into the pET28 vector

DNA coding for nbALFA-T7 RNA Polymerase was ordered as 2 IDT gBlock gene fragments, ‘pT7_6His_nbALFA_T7rnaPol’ (747-bp) and ‘T7rnaPol_T7Term’ (2600-bp) (see Table S1 and S2). These 2 gBlocks, along with the 2 pET28 amplicons of lengths 2823-bp and 2260-bp previously generated as described above, were combined to construct the plasmid using 4-fragment Gibson assembly. Multiple colonies were Sanger sequenced to identify clones with correct assembly and sequence.

### Protein purification of nbALFA-T7-RNA-Polymerase from *E. coli*

Rosetta 2 (DE3) *E. coli* cells were transformed with a sequence-verified pET28 plasmid encoding N-terminal 6xHis-tagged nbALFA-T7-RNA-Polymerase. Small scale cultures were used to optimize induction conditions, determining 18°C overnight induction with 1 mM IPTG maximized soluble nbALFA-T7-RNA-Polymerase. Rosetta 2 (DE3) *E. coli* cells from a 500 mL culture were harvested, resuspended in binding buffer (10 mM sodium phosphate, pH 7.4, 150 mM NaCl, 0.5 mM BME, Triton-X 0.025%) supplemented with protease inhibitors, and lysed by sonication. The clarified lysate was incubated with Ni-NTA agarose resin pre-equilibrated in binding buffer for 1 hour at 4°C to allow binding of 6xHis-nbALFA-T7-RNA-Polymerase. The resin was washed 3 times with 20-25 mL binding buffer supplemented with 20 mM imidazole before eluting nbALFA-T7-RNA-Polymerase with binding buffer supplemented with 500 mM imidazole. The eluate was analyzed on SDS-PAGE gels to determine purity. After concentrating the eluate with Amicon Ultra centrifugal filters, purified nbALFA-T7-RNA-Polymerase was aliquoted and stored at −80°C in binding buffer with 50% glycerol. To determine RNA polymerase activity, purified nbALFA-T7-RNA-Polymerase was intubated with rNTPs and a T7 promoter driven DNA template at 37°C for 4 hrs. Robust synthesis of expected RNA product was comparable to commercially available T7 RNA polymerase (NEB).

### Construction of small-scale PADIT-seq reporter library

We designed an oligo pool (n = 14), where each oligo contained a 3-bp randomized region, resulting in 64 unique DNA sequences per oligo that served as candidate transcription factor binding sites (TFBS). The oligo pool was obtained from IDT, mixed with an Ultramer (‘20bpBC_Bottom’) containing all possible 20-bp DNA sequences as barcodes (BCs), and double stranded using 1 cycle of KAPA HiFi polymerase. The pGL4.23 plasmid vector backbone was PCR amplified in 2 steps with Q5 High-Fidelity 2X Master Mix. First, the backbone was amplified with primers ‘pGL4.23_FWD’ and ‘pGL4.23_REV’ to exclude the luciferase open reading frame. The resulting amplicon (2359-bp) was then further amplified with primers ‘T7Term_pGL4.23_FWD’ and ‘pGL4.23_REV’ to add a 48-bp T7-terminator DNA sequence as an overlapping region for Gibson Assembly, which was performed with the resulting amplicon (2407-bp) and the double-stranded oligo pool mixed in equimolar ratios. Following chemical transformation of OneShot TOP10 cells (n = 8), ∼150,150 colonies were obtained, equivalent to ∼167 barcodes per TFBS. After recovery, and 7.5 hours of growth, the cells were maxi-prepped to obtain the small-scale PADIT-seq reporter plasmid library. Correct library assembly was validated by diagnostic PCR and Sanger sequencing of colonies.

### Obtaining TFBS-BC pairings in the small-scale PADIT-seq reporter library

To obtain TFBS-BC pairings, the library was PCR amplified using KAPA HiFi polymerase with primers, ‘AmpEZ_pGL4.2_Rev_RC’ and ‘#19_MPRA_cDNA_3.0’, which also added partial Illumina adaptors to the amplicon. Final Illumina adaptors and sample barcodes were added at Azenta, where paired-end Illumina sequencing was performed using their ‘AmpliconEZ’ service. The sequencing data was processed using custom scripts to extract TFBS-BC pairings by matching constant flanking regions. Barcodes associated with multiple TFBS were filtered out to obtain one-to-one TFBS-BC pairings.

### Construction of the all-10mers PADIT-seq reporter library

We designed and ordered two IDT Ultramers - one containing all possible 10-bp DNA sequences as candidate TFBS (‘All10mersTFBS_Top’), and another containing all possible 25-bp DNA sequences to serve as barcodes (‘25bpsBC_Bottom’). The two Ultramers were mixed in an equimolar ratio and double stranded in a single PCR cycle using KAPA HiFi polymerase. The pGL4.23 plasmid vector backbone was again PCR amplified in 2 steps with Q5 High-Fidelity 2X Master Mix. First, the backbone was amplified with primers ‘pGL4.23_FWD’ and ‘pGL4.23_REV’ to exclude the luciferase open reading frame. The resulting amplicon (2359-bp) was then further amplified with primers ‘T7Term_pGL4.23_F_2.0’ and ‘pGL4.23_REV’ to add a 56-bp DNA sequence as an overlapping region for Gibson Assembly, which was performed with the resulting amplicon (2415-bp) and the double-stranded oligo-pool mixed in equimolar ratios. Following desalting with a mixed cellulose esters (MCE) hydrophilic membrane (0.025 um), the assembled reporter library plasmid was electroporated into E. cloni 10G Supreme cells (n = 13 transformations). Based on plating experiments, the total number of transformants obtained was estimated to be 110 million, providing an average of ∼100 barcodes per TFBS. The transformed cells were recovered, grown for 6.5 hours, and maxi-prepped to obtain the complete all-10mers PADIT-seq reporter plasmid library containing over 100 million clones. Correct library assembly was validated by diagnostic PCR and Sanger sequencing of 10 colonies.

### Obtaining TFBS-BC pairing in the all-10mers PADIT-seq reporter library

To obtain TFBS-BC pairings, the all-10mers PADIT-seq reporter library was PCR amplified using KAPA HiFi polymerase. Four forward primers were designed with partial Illumina adapters, 6N randomized bases, and 2-bp staggers (‘All_10mers_LibSeq_F1-4’). These were used individually with a single reverse primer (‘All_10mers_LibSeq_R’) to generate 4 PCR-1 products of expected sizes 213-219-bp (9 cycles). The 4 PCR-1 products were then used as template in PCR-2 (5 cycles) with TruSeq indexed primers to attach Illumina sample indexes. This generated 4 PCR-2 products of expected size 272-278-bp. After PCR amplification, the 4 products were SPRI cleaned and analyzed on an Agilent TapeStation to confirm expected sizes. The 4 indexed libraries were sequenced separately on a NovaSeq6000 (2×150 bp reads). The sequencing data from each of the 4 indexed libraries (F1-F4) was combined and processed using custom scripts to extract unique TFBS-BC combinations along with their counts by matching the constant flanking regions. Barcodes unambiguously associated with only 1 TFBS across all 4 libraries were classified as ‘single TFBS barcodes’ and retained.

The all-10mers PADIT-seq reporter plasmid library was amplified in 4 separate PCR reactions (F1-F4) with different TruSeq indexes to identify potential PCR-mediated recombination artifacts in the following way: for barcodes associated with multiple TFBS, an initial filter retained only TFBS observed independently in at least 2 of the 4 libraries. The rationale being that TFBS-BC occurrence in multiple independent libraries indicates likely true pairings versus artifacts of PCR-mediated recombination. After this first multi-library filtering step, any barcodes still associated with multiple TFBS were removed entirely to eliminate ambiguities. As an additional filter, barcodes where the top TFBS had fewer reads than the sum of discarded TFBS were removed. The ‘single TFBS barcodes’ and vetted multiple TFBS barcodes were combined to obtain high-confidence 1:1 TFBS-BC pairs for downstream analysis. This multi-step filtering process leveraged the independently prepared sequencing libraries to remove incorrect and ambiguous TFBS-BC pairings arising from PCR-mediated recombination. It enabled retaining high-confidence barcode-TFBS pairs reproducibly identified across multiple libraries while discarding likely PCR artifacts and errors.

### PADIT-seq experiments

To remove any supercoiling, PADIT-seq reporter libraries were first linearized with DrdI (NEB), which cuts a 12-bp DNA sequence (GACNNNN/NNGTC) only once in the pGL4.23 vector. For every DBD being tested, the following 30 ul PURExpress IVTT reactions (NEB) were assembled: 10 ul Solution A, 7.5 ul Solution B, 1 ul Murine RNase Inhibitor, 3 ul 100 mM rNTPs, 0.45 ul 1000 mM magnesium acetate, 3 ul previously purified nbALFA-T7-RNA-Polymerase, ∼300 ng linearized PADIT-seq reporter plasmid library, ‘pT7-ALFA-DBD-T7Term’. The linearized PADIT-seq reporter plasmid library was mixed with ‘pT7-ALFA-DBD-T7Term’ amplicons in an approximately 2:1 molar ratio. The 30 ul PURExpress IVTT reactions were split into three wells, and all subsequent steps were performed separately (3 biological replicates).

### cDNA synthesis of PADIT-seq reporter RNAs

After 4 hours at 37°C, the 10 ul reactions were purified with RNAClean XP as per manufacturer’s instructions, eluting in 35 ul Nuclease-free water. 2 ul barcoded cDNA synthesis primers (0.1uM final each) were added to 18 ul purified RNA, incubated at 75°C for 3 mins, then placed on ice. cDNA was synthesized by adding 10 2X Multiscribe reaction mix (Thermo Fisher), and incubating at 25°C for 20 minutes, followed by 37°C for 120 minutes. Minus reverse transcriptase controls were performed in parallel. Excess primers were removed from the cDNA:RNA duplexes by adding Exonuclease I and incubating at 37°C for 60 mins, followed by heat inactivation at 80°C for 20 mins. Quantitative PCR was performed to verify degradation of all excess primers, and to determine the threshold cycle of sample cDNAs.

### PADIT-seq library preparation for Illumina sequencing

For the small-scale PADIT-seq library, barcoded cDNAs synthesized from the reporter RNAs were pooled prior to PCR amplification. The pooled cDNA was amplified in a single PCR reaction using KAPA HiFi polymerase with primers ‘MPRA_AmpliconEZ_FWD’ and ‘MPRA_AmpEZ_REV2.0’. This generated a PCR-1 product that was then used as template for a second PCR with primers ‘#34_MPRA’ and ‘169_TruSeq_Multiplex_220_2’ to attach Illumina adapters and sample barcodes. In contrast, for the all-10mers PADIT-seq library, barcoded cDNAs were kept separate and amplified in individual PCR reactions rather than pooled. For each DBD, cDNA was amplified using KAPA HiFi HotStart Polymerase with primers ‘MPRA_AmpliconEZ_FWD’ and ‘MPRA_AmpEZ_REV2.0’. This generated PCR-1 products that were cleaned and quantified. In the second PCR, ‘#34_MPRA’ and indexed TruSeq primers were used to attach Illumina adapters and sample barcodes. Aiming for 50X coverage, Illumina sequencing was performed on NovaSeq 6000 (sample sequencing statistics in Table S4).

### PADIT-seq differential activity analysis

Barcodes from sequencing libraries were mapped to the associated TFBS based on previously obtained TFBS-BC pairings. Barcode counts per TFBS were obtained for each library and merged into a single data frame. Quality control was performed by generating Pearson correlation heatmaps and principal component analysis (PCA) plots to assess reproducibility between replicates and overall structure of the data. For differential activity analysis, read counts for the DBD-of-interest and a ‘no DBD’ control, across 3 replicates each, were analyzed using DESeq2. TFBS significantly bound by the DBD-of-interest were identified by applying a false discovery rate (FDR) threshold of 5%.

### Extracting 8-mer and 9-mer PADIT-seq activities from the all-10mers PADIT-seq library

To extract HOXD13 8-mer PADIT-seq log_2_ fold change values, all possible occurrences of each 8-mer in the all-10-mers PADIT-seq library were analyzed. This was done separately for 8-mers at the 3 different positions within the 10-mers, where median log_2_ fold change values were retained for each position. Both forward and reverse orientations were analyzed and the orientation with the higher median log_2_ fold change was retained for each position within the 10-mers. The median values across the 3 positions were then compared to HOXD13 PBM E-scores (Table S5). To extract EGR1 9-mer PADIT-seq log_2_ fold change values, all possible occurrences of each 9-mer in the all-10-mers PADIT-seq library were analyzed. This was done separately for 9-mers at the 2 different positions within the 10-mers, where median log_2_ fold change values were retained for each position. Both forward and reverse orientations were analyzed and the orientation with the higher median log_2_ fold change was retained for each position within the 10-mers. The median values across the 2 positions were then compared to EGR1 PBM E-scores (Table S6). Area under the receiver operating characteristic (AUROC) for the ability of PBM E-scores to classify PADIT-seq active *k*-mers were >0.985 for both TFs. The PBM E-score thresholds, 0.30 and 0.45 for HOXD13 and EGR1, respectively, correctly classified 70% of the PADIT-seq active *k*-mers with <1% false positives.

### Comparing PADIT-seq activities between the all-10mers and small-scale libraries

For each 9-bp TFBS tested in the smaller focused PADIT-seq library (n = 896), the median log_2_ fold change value was extracted from the 8 possible occurrences of the 9-bp TFBS in the all-10mers PADIT-seq library.

### EGR1 9-mer E-scores

EGR1 PBM data from UniPROBE was re-analyzed using the Universal PBM Analysis Suite modified to obtain 9-mer instead of 8-mer E-scores (*3, 4*).

### Analysis of HT-SELEX data

Publicly available HOXD13 and EGR1 HT-SELEX sequencing data (*45*) were downloaded and analyzed with default parameters using the R SELEX package (*7, 81*). *K*-mers were enriched if the observed to expected counts were greater than 3.

### Obtaining ProBound relative affinities

MotifCentral PWM model numbers 17019 and 12718 were used for HOXD13 and EGR1, respectively. Because both PWM models were > 9-bp, the *k*-mers are padded with N’s to match the length of PWM models. ProBound relative affinity for all possible permutations of N padding around a *k*-mer were calculated and the highest affinity values were retained for all 8-mers (HOXD13) and 9-mers (EGR1).

### Analysis of ChIP-seq data

Hoxd13 and Egr1 ChIP-seq data were processed using the ReMap database pipeline (*82*). Briefly, adapters were removed using Trim Galore, trimming reads up to 30-bp, and Bowtie2 (*83*) with options -end-to-end and -sensitive was used to align all reads on the mouse genome (mm10). Hoxd13 and Egr1 ChIP-seq peak calls were downloaded from the ReMap database. Random length-matched genomic intervals were generated using the ‘shuffle’ command in Bedtools (*84*). Sequence logos were generated using R package ‘seqLogo’.

### Evolutionary conservation analysis

PhastCons scores (*85*) were downloaded in BigWig format from the UCSC genome browser (‘mm10.60way.phastCons60wayGlire.bw’). For every genomic region of interest, PhastCons scores were mapped to each nucleotide within using the Bedtools map command (*84*). Genomic regions with missing values were filtered out, and paired Wilcoxon tests were performed to statistically compare PhastCons scores of consecutive nucleotides.

### Network analysis

Custom R scripts (see code availability) were written to obtain adjacency matrices from *k*-mer DNA sequences using the logic described in the main text. The fraction of *k*-mers within the largest, single connected component was obtained using ‘igraph’ R package. For network analysis using PBM data, only TFs with top 500 8-mers with E-score > 0.35 were analyzed.

### Differential PADIT-seq activity between variant alleles

Variants were predicted to alter TF binding if two conditions were met: 1) the total number of active overlapping *k*-mers in the two alleles were not identical, and 2) the absolute difference in the sum of PADIT-seq activities across both the alleles was >1. The second condition applies a minimum threshold on the effect size of variants effects on TF binding, i.e., if the variant were to be tested by PADIT-seq, we would observe a >2-fold change in reporter gene expression between the two alleles.

---

**Figure SI:**
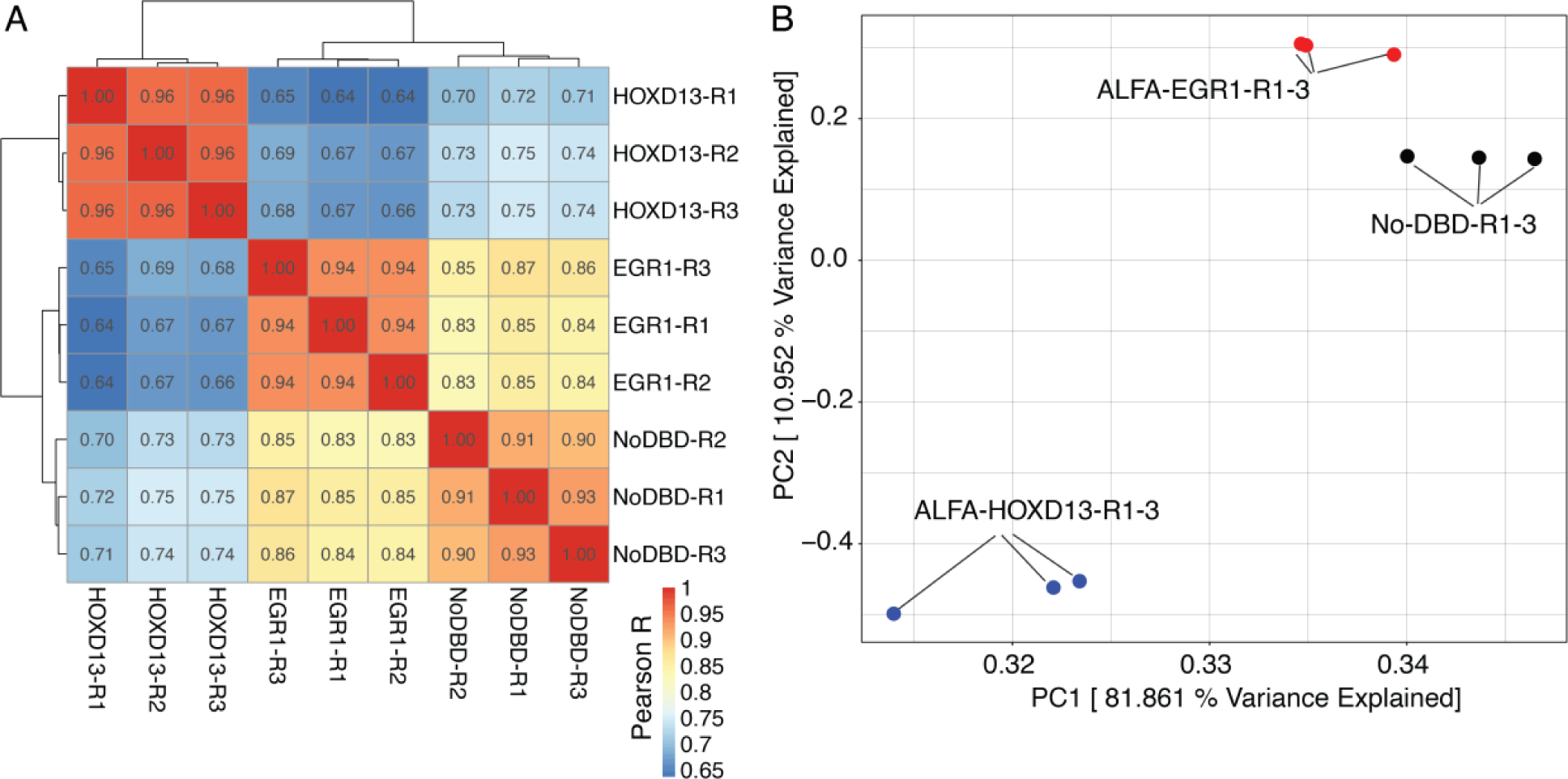
All-IOmer TFBS counts are highly reproducible across triplicate PADIT-seq assays. **(A)** Heatmap of pairwise Pearson correlation coefficients with unsupervised row and column clustering. R1-3 denotes replicates. **(B)** Scatter plot of the first two principal components, which together explains ∼93% of the variation in the 9 PADIT-seq libraries.

**Figure S2:**
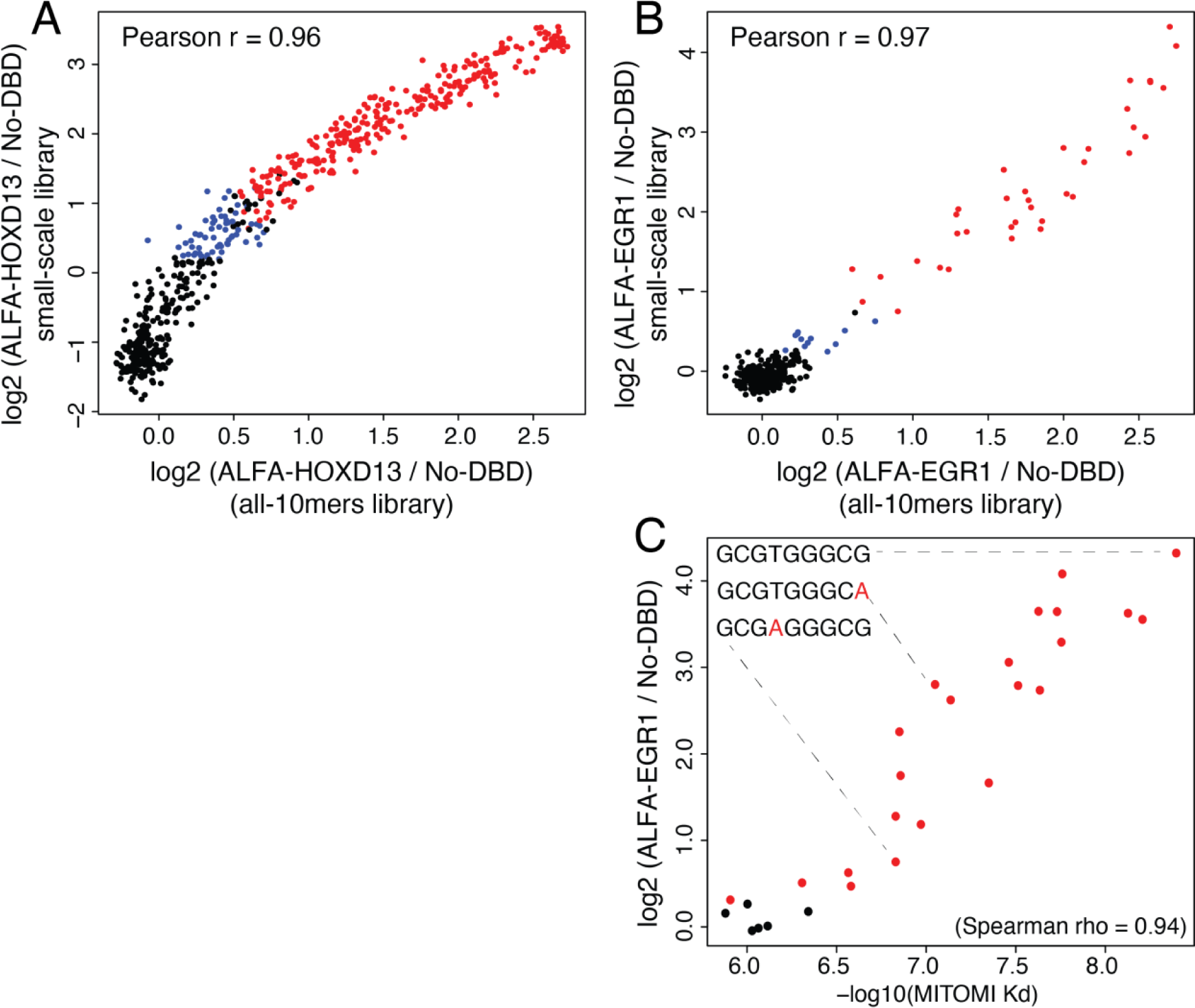
PADIT-seq activity from the all-10mers and small-scale library are highly correlated. **(A-B)** PADIT-seq activities for HOXD13 (A) and EGR1 (B) from the all-1 Omers library and the small-scale library are compared. Red TFBS are active in both libraries. Black TFBS are not active in either libraries. Blue TFBS are active only in the small-scale library. **(C)** PADIT-seq activity from the small-scale library and MITOMI-derived dissociation constants (Kd) for EGR1 are compared. Red TFBS are active, whereas black TFBS are not active.

**Figure S3:**
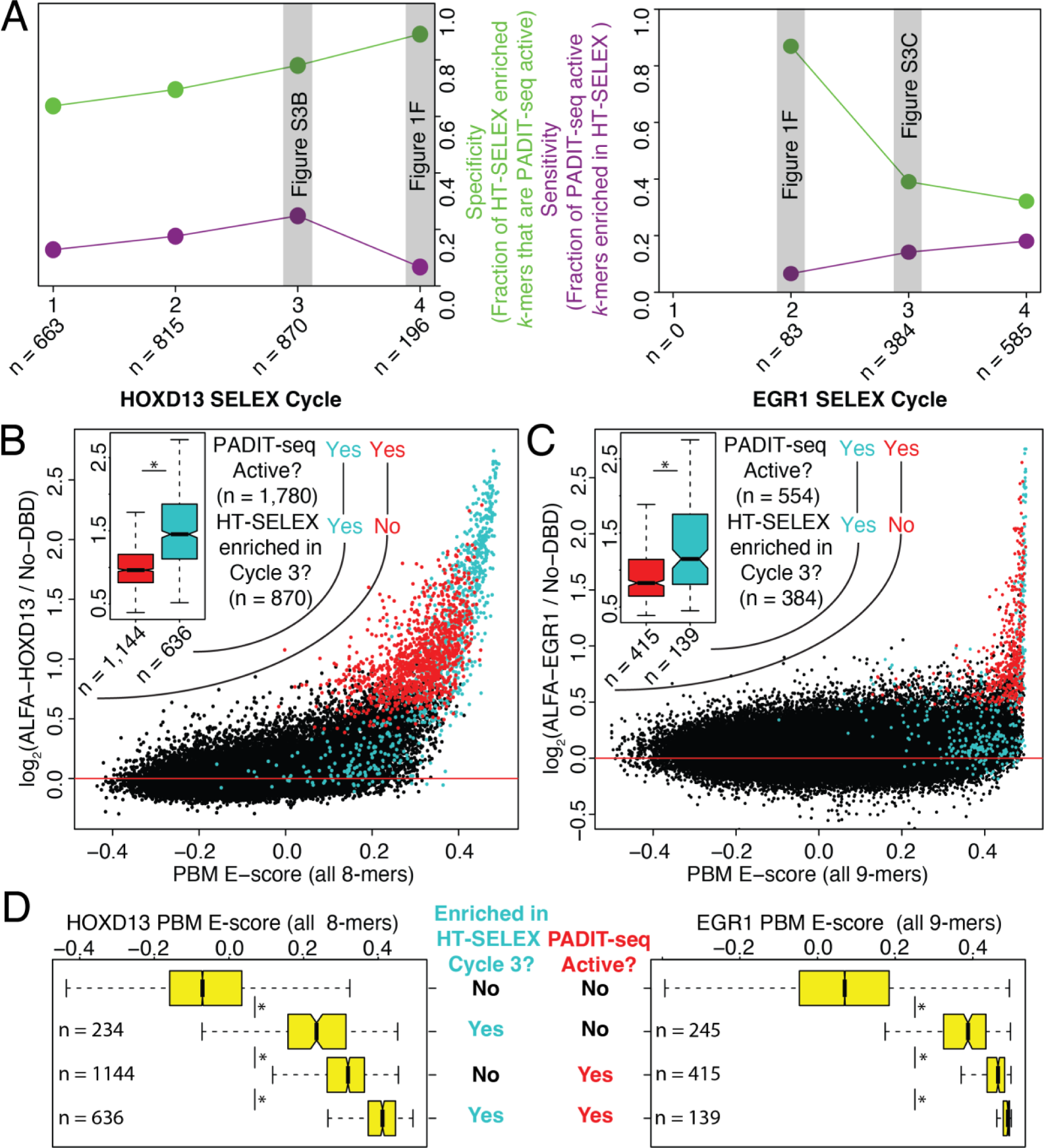
HT-SELEX enriched ƙ-mers are biased for detecting high affinity TFBS. **(A)** Specificity and sensitivity of HT-SELEX cycles 1-4 in detecting PADIT-seq active ƙ-mers for H0XD13 (left) and EGR1 (right). Specificity is defined as the fraction of HT-SELEX enriched ƙ-mers that are PADIT-seq active. Sensitivity is defined as the fraction of PADIT-seq active ƙ-mers that are enriched in HT-SELEX. **(B-C)** Plots in Figure 1E are shown again with points colored differently. Blue TFBS are significantly enriched in HT-SELEX cycle 3. **(inset)** PADIT-seq activities for active TFBS enriched (blue) or not enriched (red) in HT-SELEX are compared. * denotes a Wilcoxon rank sum test p-value < 0.05. **(D)** All 8-mers for HOXD13 (left) and 9-mers for EGR1 (right) were categorized based on whether they were enriched in HT-SELEX cycle 3 and whether they were PADIT-seq active. Corresponding PBM E-scores are plotted on the y-axis. * denotes Wilcoxon test p-value < 2.2 x 10^16^.

**Figure S4:**
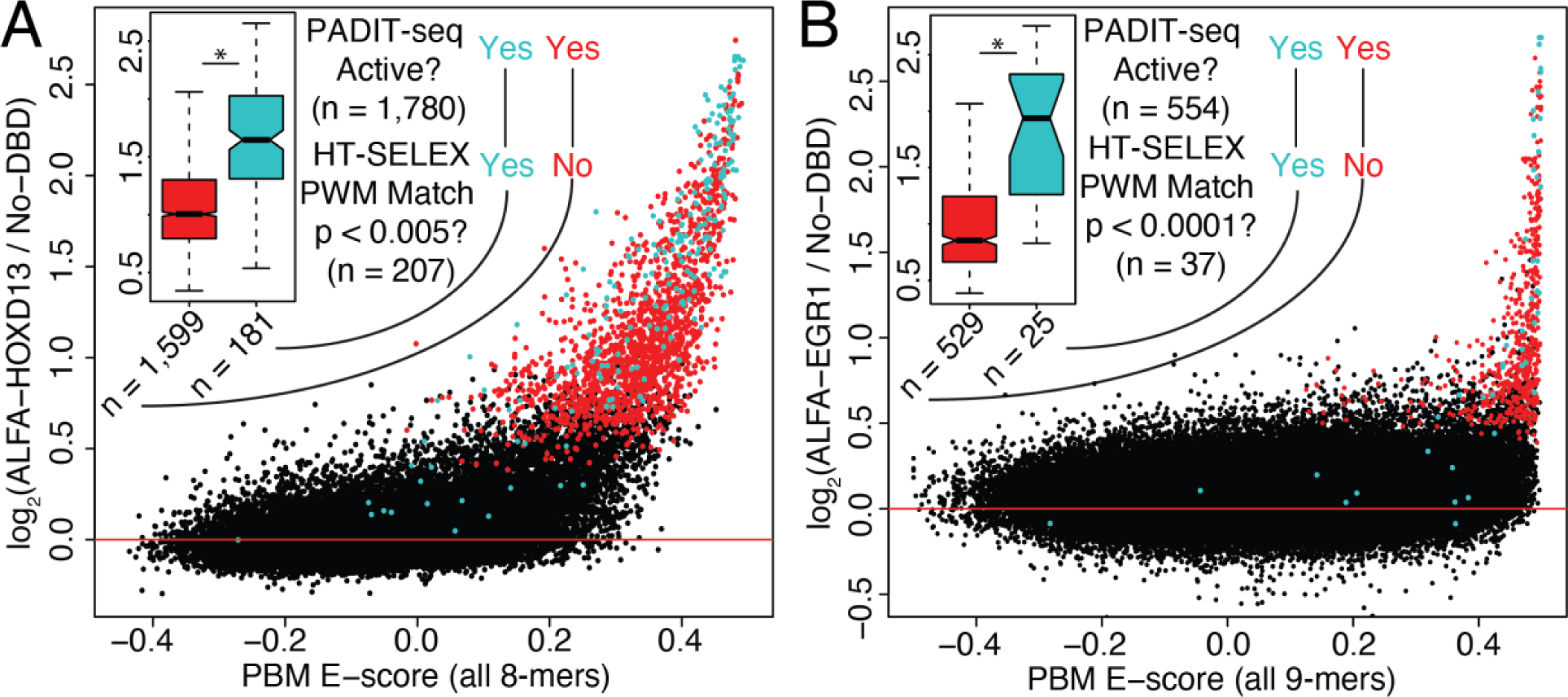
PWMs derived from HT-SELEX data are biased for detecting high affinity TFBS. **(A-B)** Plots in Figure 1E are shown again with points colored differently. Blue TFBS are a significant PWM match (FIMO). PWMs were derived from HT-SELEX data, **(inset)** PADIT-seq log_2_(fold change) values for active TFBS that are a significant PWM match (blue) or not (red) are compared. * denotes a Wilcoxon rank sum test p-value < 0.05.

**Figure S5:**
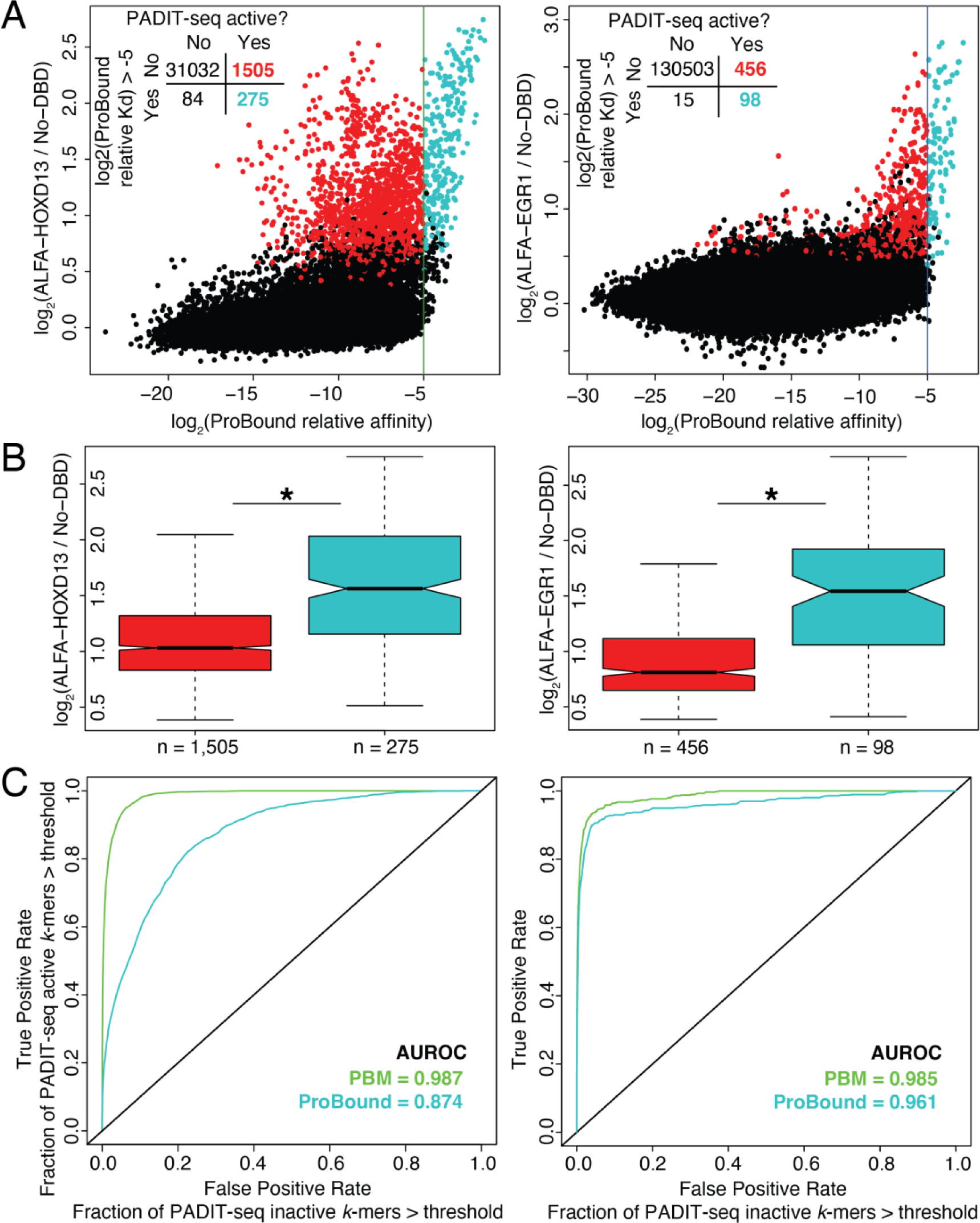
ProBound relative affinities are biased for detecting high affinity TFBS. **(A)** PA-DIT-seq activities (y-axis) are compared to ProBound-derived relative affinities for all 8-mers (HOXD13, left) and 9-mers (EGR1, right) derived from HT-SELEX PWM model numbers 17019 and 12718, respectively, in the MotifCentral database. Black points are PADIT-seq inactive ƙ-mers, red are active ƙ-mers below the ProBound relative affinity threshold and blue are active ƙ-mers above the ProBound relative affinity threshold. **(B)** PADIT-seq log_2_(fold change) values for active TFBS with log_2_(ProBound relative affinity) greater than (blue) or less than (red) −5 are compared. * denotes a Wilcoxon rank sum test p-value < 0.05. **(C)** Fraction of PADIT-seq active ƙ-mers above classification threshold (True Positive Rate) versus the fraction of PADIT-seq inactive k-mers above classification threshold (False Positive Rate) for HOXD13 (left) and EGR1 (right). The performance of PBM E-scores (green) and ProBound relative affinities (blue) are compared, and AUROC values are indicated.

**Figure S6:**
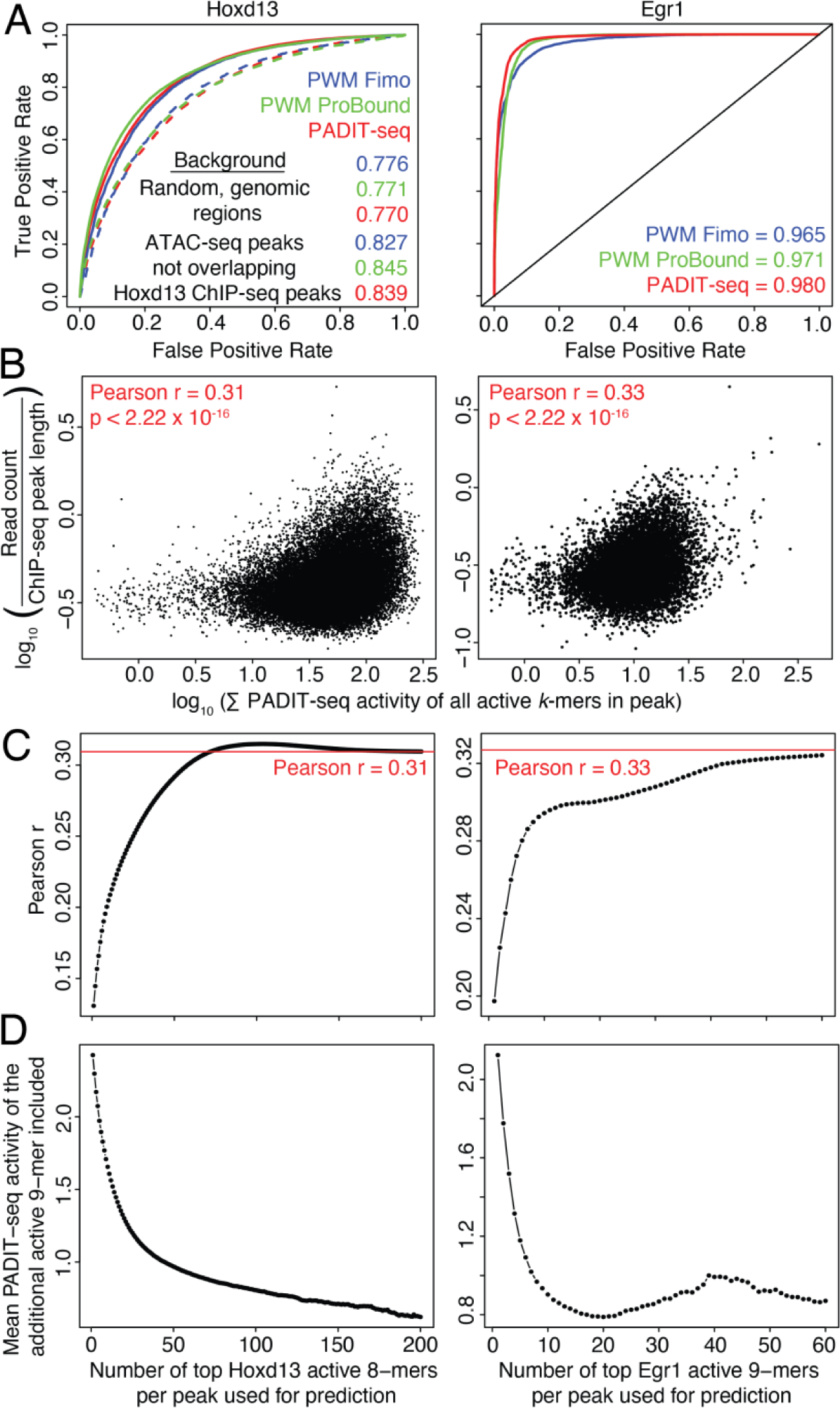
Lower affinity binding sites increase TF genomic occupancy at ChIP-seq peaks. **(A)** Frac-tion of Hoxdl 3 ChIP-seq peaks above classification threshold (True Positive Rate) versus the fraction of background genomic intervals above the same threshold (False Positive Rate) for Hoxd13 (left) and Egr1 (right). For Hoxdl 3, in addition to random, length matched background genomic intervals, false positives were also deter-mined with the background sequenc-es defined to be embryonic forelimb bud ATAC-seq peaks not overlapping Hoxdl3 ChIP-seq peaks. **(B)** The sum of PADIT-seq activities of all the active k-mers in Hoxd13 (left) and Egr1 (right) ChIP-seq peaks is plotted against the corresponding read counts normalized to peak length. **(C)** The number of active k-mers whose PADIT-seq activities are summed is varied on the x-axis. Only the specified number of top active k-mers are used to predict the normalized ChIP-seq read counts for Hoxdl3 (left) and Egr1 (right). The corre-sponding Pearson correlation coeffi-cients are plotted on the y-axis. **(D)** For every additional active k-mer per peak used to predict the normalized ChIP-seq read counts, the corresponding mean PADIT-seq activities are plotted on the y-axis.

**Figure S7:**
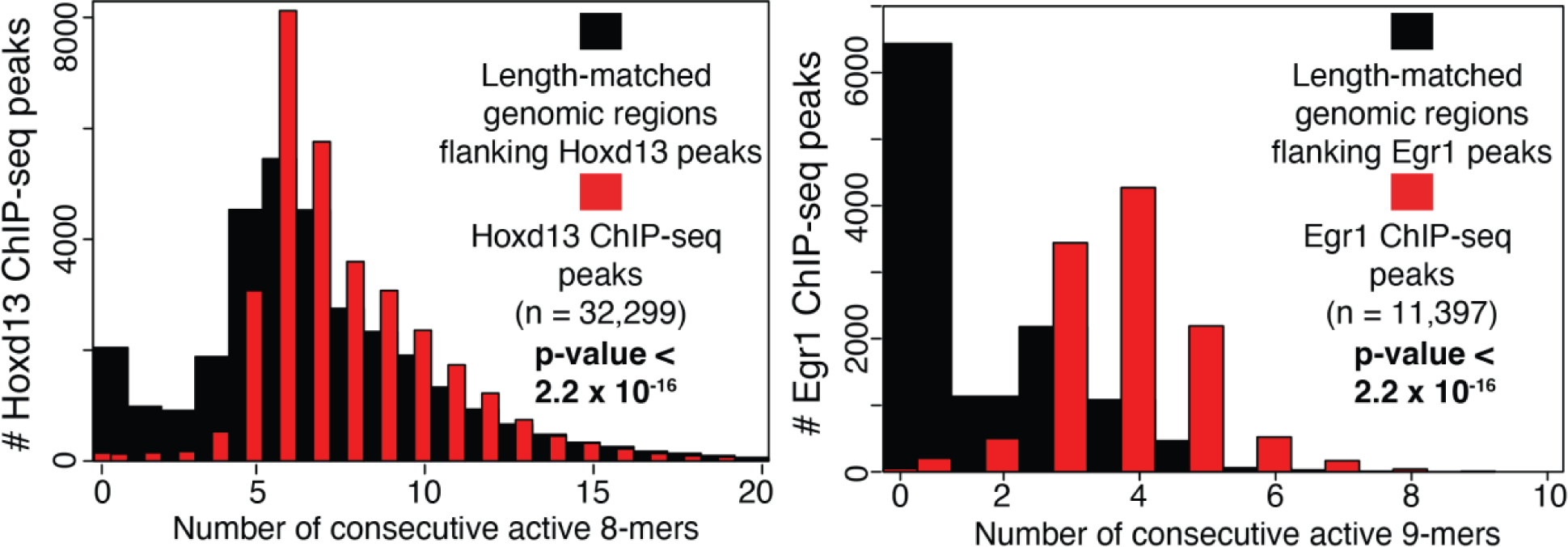
Hoxd13 and Egr1 ChIP-seq peaks have significantly more consecutive active k-mers, irrespective of how background genomic sequences were defined. Histogram of the number of consecutive active ƙ-mers in Hoxd13 (left) and Egr1 (right) ChIP-seq peaks. Background genomic intervals were generated by shifting the peak interval one peak length to the left.

**Figure S8:**
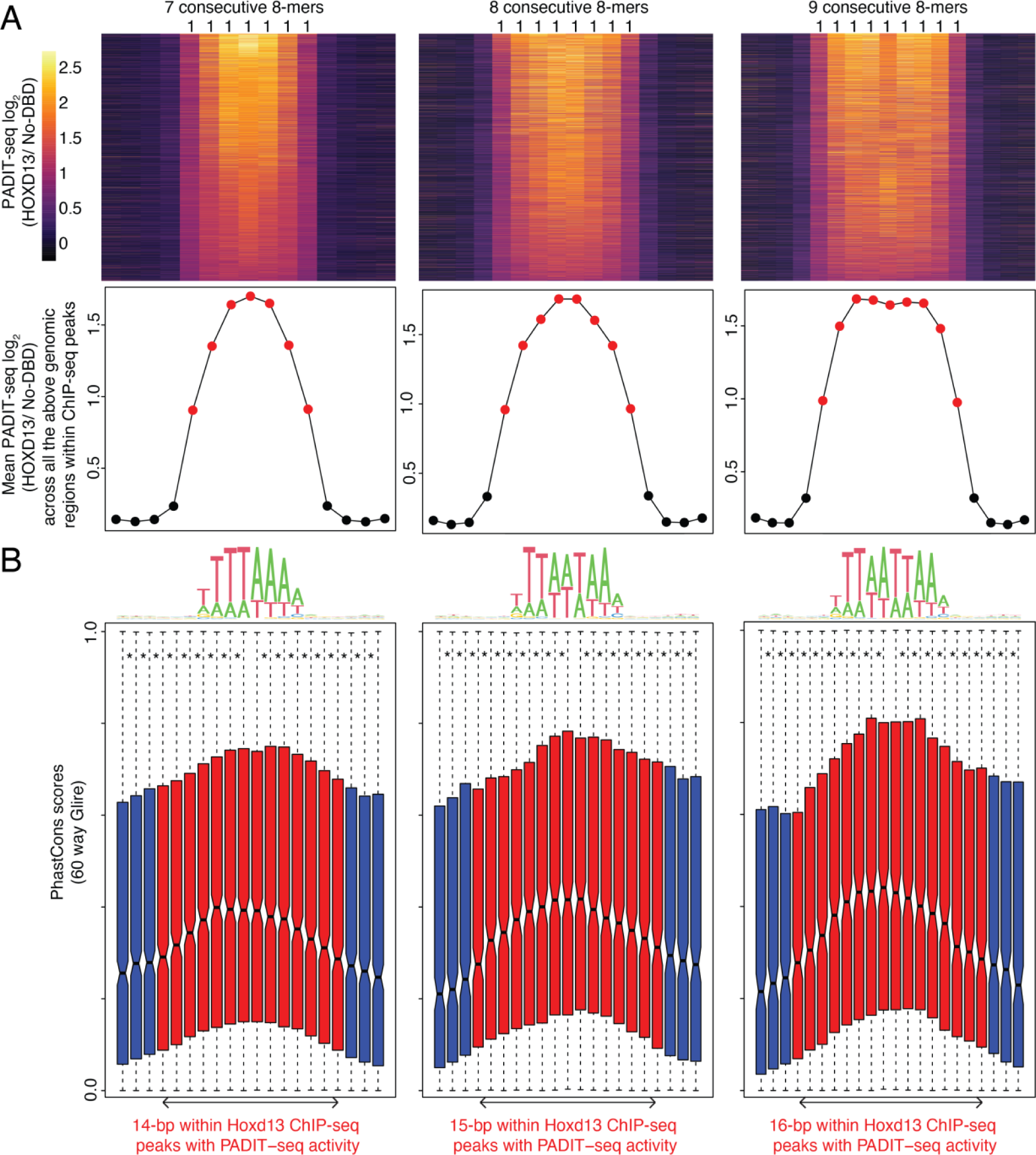
Genomic regions containing consecutive active HOXD13 k-mers are evolutionarily conserved. **(A)** (Top) Heatmap of PADIT-seq activity at Hoxd13 ChIP-seq peaks with 7-9 consecutive active 8-mers, along with 4 flanking inactive 8-mers. Each row is a ChIP-seq peak. (Bottom) Mean PADIT-seq activities of all the genomic regions within ChIP-seq peaks in the heatmap above. (B) 60-way Glire PhastConε scores for the genomic regions containing 7-9 consecutive active 8-mers in Hoxdl 3 ChIP-seq peaks (red). 60-way Glire PhastCons scores for the flanking 3 nucleotides are in blue. Adjusted p-values < 0.05 from paired Wilcoxon rank sum tests are indicated by *.

**Figure S9:**
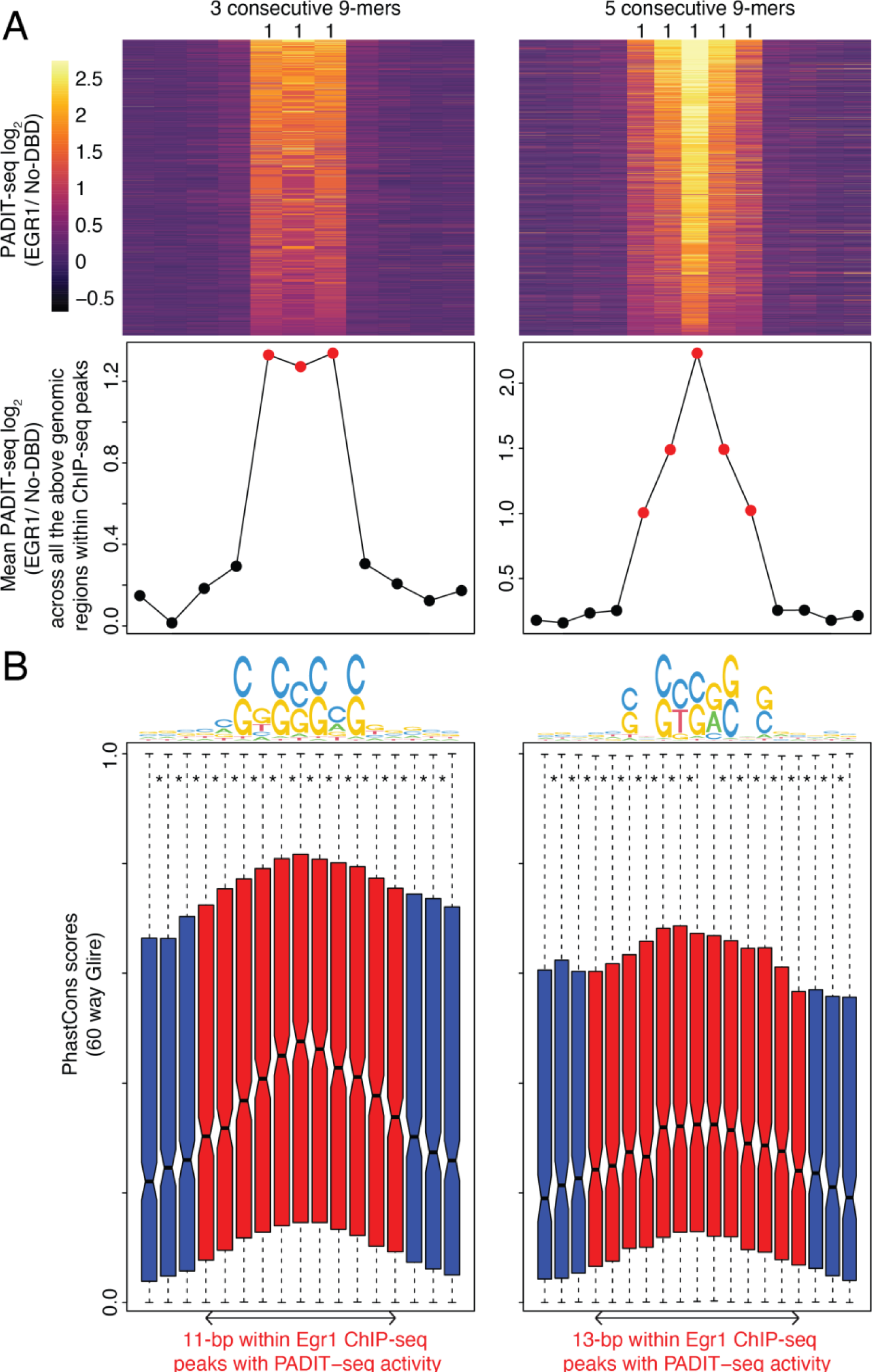
Genomic regions containing consecutive active EGR1 fr-mers are evolutionarily conserved. **(A)** (Top) Heatmap of PADIT-seq activity at Egr1 ChIP-seq peaks with 3 (left) or 5 (right) consecutive active 9-mers, along with 4 flanking inactive 9-mers. Each row is a ChIP-seq peak. (Bottom) Mean PADIT-seq activities of all the genomic regions within ChIP-seq peaks in the heatmap above. **(B)** 60-way Glire PhastCons scores for the genomic regions containing 3 (left) or 5 (right) consecutive active 9-mers in Egr1 ChIP-seq peaks (red). 60-way Glire PhastCons scores for the flanking 3 nucleotides are in blue. Adjusted p-values < 0.05 from paired Wilcoxon rank sum tests are indicated by *.

**Figure S10:**
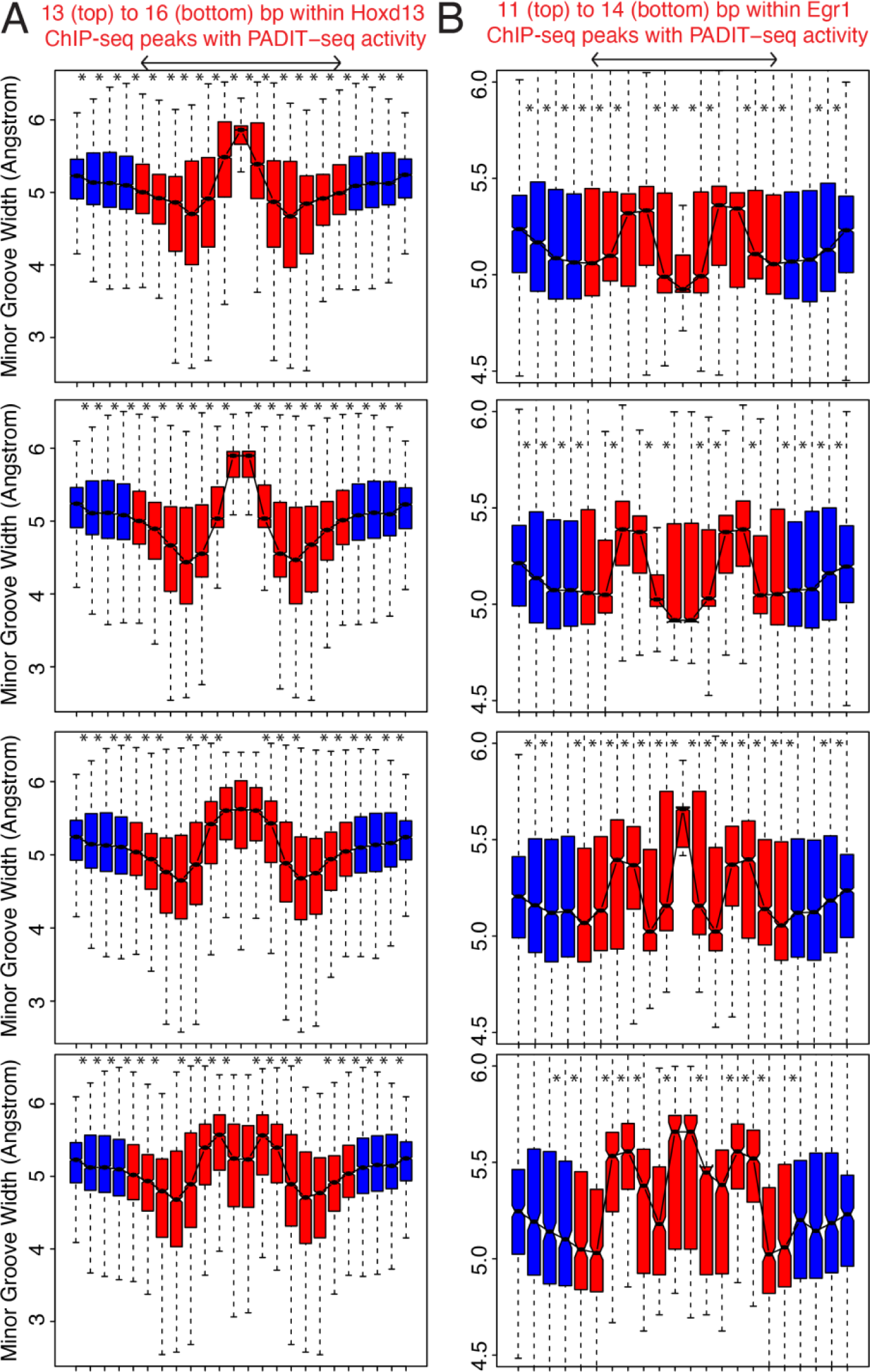
MGW at the extended recognition sequences bound by Hoxdl3 and Egr1 are distinct from flanking genomic regions. **(A)** For Hoxdl3 ChIP-seq peaks with 6-9 consecutive active 8-mers, predicted minor groove widths (MGW) for the corresponding genomic regions (red), including 4-bp flanks (blue), are plotted on the y-axis. **(B)** For Egrl ChIP-seq peaks with 3-6 consecutive active 9-mers, predicted minor groove widths (MGW) tor the corresponding genomic regions (red), including 4-bp flanks (blue), are plotted on the y-axis. **(A-B)** Per base MGW values for every genomic region in ChIP-seq peaks were predicted using ‘Deep DNAShape’. Adjusted p-values < 0.05 from paired Wilcoxon rank sum tests are indicated by *.

**Figure S11:**
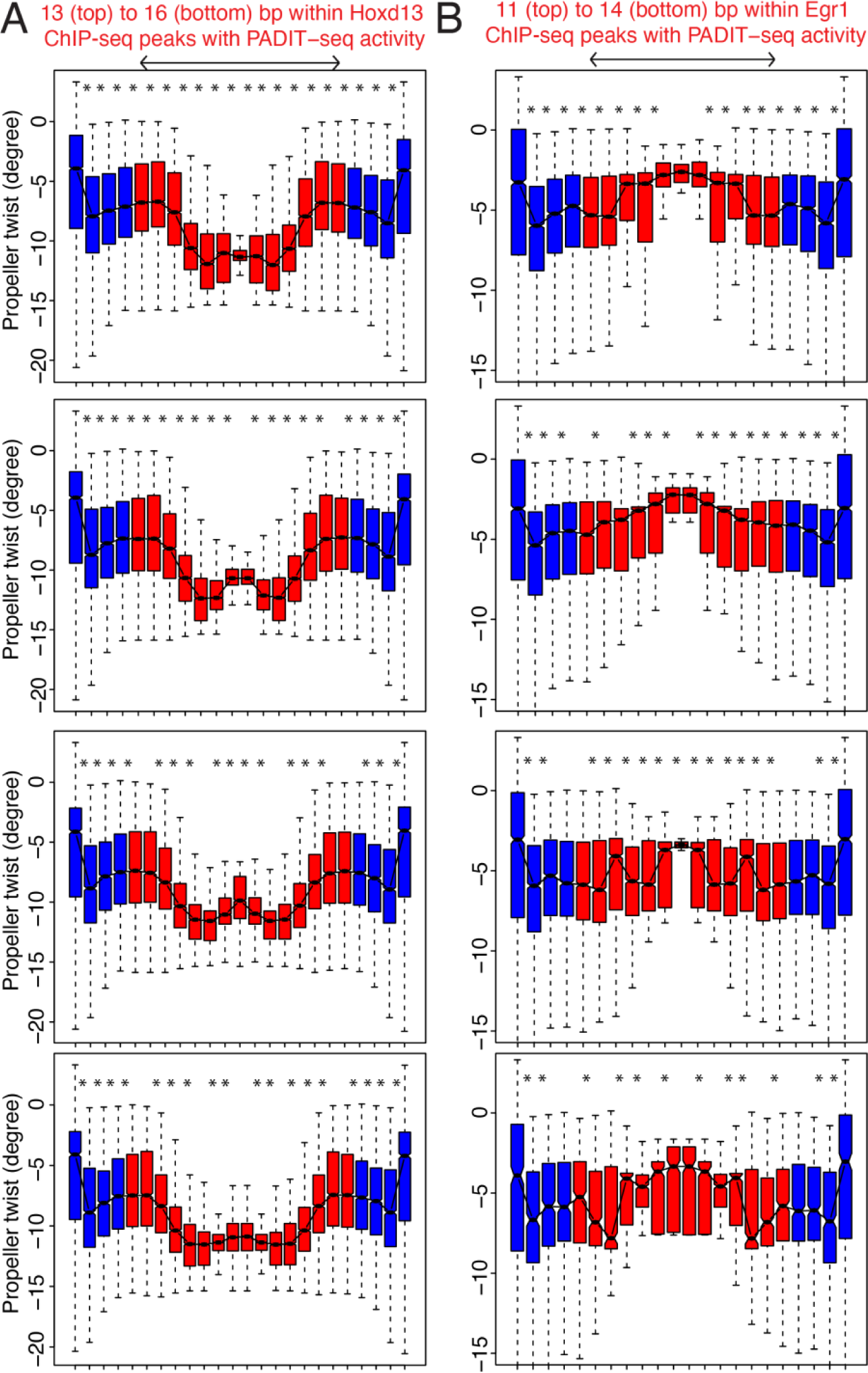
Propeller twist at the extended recognition sequences bound by Hoxdl3 and Egr1 are distinct from flanking genomic regions. **(A)** For Hoxdl3 ChIP-seq peaks with 6-9 consecutive active 8-mers, predicted propeller twist (ProT) for the corresponding genomic regions (red), including 4-bp flanks (blue), are plotted on the y-axis. **(B)** For Egr1 ChIP-seq peaks with 3-6 consecutive active 9-mers, predicted ProT for the corresponding genomic regions (red), including 4-bp flanks (blue), are plotted on the y-axis. **(A-B)** Per base ProT values for every genomic region in ChIP-seq peaks were predicted using ‘Deep DNAShape’. Adjusted p-values < 0.05 from paired Wilcoxon rank sum tests are indicated by *.

**Figure S12:**
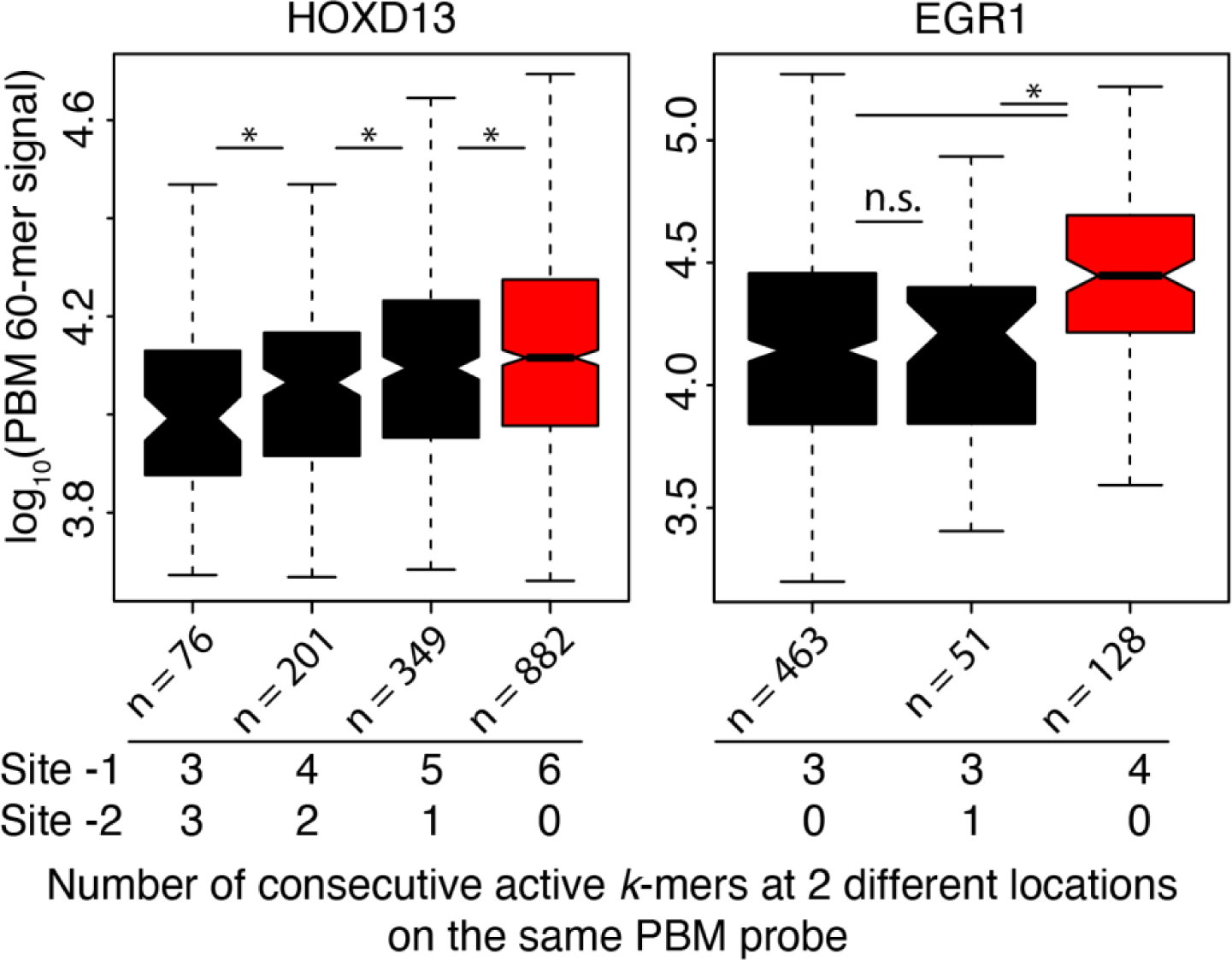
Overlapping binding sites increase TF occupancy to a higher degree than equivalent numbers of non-overlapping binding sites. PBM 60-bp probe signal intensity as a function of the number of consecutive overlapping active ƙ-mers at 2 distinct non-overlapping sites on the same probe. * indicates a Wilcoxon test adjusted p-value < 0.05.

